# Undecaprenyl Pyrophosphate Phosphatase (UppP) is a pivotal element in *Salmonella* Intramacrophage Survival

**DOI:** 10.1101/2025.04.24.650467

**Authors:** Rhea Vij, Debapriya Mukherjee, Kirti Parmar, Krishna Chaitanya Nallamotu, Manjula Reddy, Dipshikha Chakravortty

## Abstract

To establish infection, *Salmonella* confronts a dynamic barrage of host-induced stresses. The peptidoglycan layer, essential for maintaining cell integrity and counteracting environmental stress, relies on the lipid carrier undecaprenyl phosphate, generated by Undecaprenyl pyrophosphate phosphatase (UppP). While UppP is linked to virulence in other pathogens, its role in *Salmonella* remains unclear. We show that an *uppP* mutant in *S*. Typhimurium exhibits altered cell morphology, reduced stiffness, and impaired survival in RAW 264.7 macrophages. The mutant is also attenuated in systemic infection in C57BL/6 mice. These defects are associated with increased sensitivity to nitrosative stress. Notably, iNOS inhibition or deficiency restores intracellular survival of the *uppP* mutant in both RAW 264.7 macrophages and the mouse model, implicating UppP in resistance to nitrosative stress. Our findings reveal a critical role for UppP in promoting *Salmonella* survival within macrophages and contributing to systemic pathogenesis.

## Introduction

*Salmonella* is a Gram-negative foodborne pathogen belonging to the *Enterobacteriaceae* family [1]. The genus, comprising more than 2600 serovars, can infect various hosts, including plants, birds, and mammals[2], [3]. Clinical manifestations of *Salmonella* infection range from self-limiting gastroenteritis associated with non-typhoidal serovars to systemic infection caused by typhoidal serovars and invasive non-typhoidal serovars [4]. The significant burden of *Salmonella*-associated enterocolitis, affecting 95.1 million people in 2017, coupled with the rising threat of drug-resistant strains as recognised by its inclusion on the WHO priority pathogen list, underscores *Salmonella*’s growing danger to public health [5], [6], [7].

*Salmonella* enters the host through contaminated food and water. *Salmonella* encounters several stresses within the host, including oxidative and nitrosative stress, osmotic stress, acidic pH, nutrient starvation, metal ions, and host antimicrobials [8]. Out of all these stresses, the bacterial peptidoglycan layer helps counter osmotic stress by acting as a barrier to the external environment and resisting internal turgor pressure, thereby preventing the cell from bursting. It also helps in maintaining the bacterial shape and virulence. The synthesis of the peptidoglycan layer is a concerted effort of many proteins and takes place in the cytosolic and periplasmic compartments of the bacterial cell [9]. The biosynthesis of the peptidoglycan building block, N-acetylglucosamine-N-acetyl muramyl-pentapeptide, takes place in the cytoplasm on the lipid carrier undecaprenyl phosphate (C55-P), forming Lipid II. Lipid II is then flipped by MurJ to the periplasmic phase [9], [10]. In the periplasmic phase, the disaccharide pentapeptide part of Lipid-II undergoes polymerisation by SEDS (Shape, Elongation, Division, and Sporulation) proteins (such as FtsW and RodA) and Penicillin-binding proteins (PBPs) for peptidoglycan synthesis, releasing undecaprenyl pyrophosphate [11], [12], [13], [14]. The undecaprenyl pyrophosphate (C55-PP) is dephosphorylated by undecaprenyl pyrophosphate phosphatase (UppP) to undecaprenyl phosphate in the periplasmic phase, which is recycled for another round of peptidoglycan synthesis [15]. While UppP accounts for approximately 75% of this phosphatase activity in *E. coli*, the remaining activity is provided by the functionally redundant PAP2 family proteins, PgpB, YbjG, and LpxT [16], [17].

Certain studies have highlighted the crucial role of undecaprenyl pyrophosphate phosphatases in the virulence and pathogenesis of different bacterial species. In *E. coli,* the undecaprenyl pyrophosphate phosphatase encoded by the gene *uppP*, also known as *bacA*, leads to bacitracin resistance when it is overexpressed. Bacitracin strongly binds to undecaprenyl pyrophosphate and prevents its dephosphorylation [18]. Mutant strains lacking the *bacA* homologue in *Staphylococcus aureus* and *Streptococcus pneumoniae* exhibited reduced virulence and severe attenuation, respectively, in the mouse model of infection [19]. HupA, an undecaprenyl pyrophosphate phosphatase of the PAP2 family, belonging to *Helicobacter pylori*, is essential for colonisation of the stomach [20]. Other reports have shown that deletion of *uppP* in *Mycobacterium smegmatis* impairs its ability to form biofilms [21]. However, the role of undecaprenyl phosphate in *Salmonella* Typhimurium (STM) pathogenesis has remained unexplored.

Recent studies suggest that intracellular *Salmonella* actively remodels its peptidoglycan layer to survive in the intracellular niche, differing significantly from those grown in LB medium [22], [23]. Intracellular *Salmonella* Typhimurium actively edits its peptidoglycan to incorporate elevated levels of L-alanine-D-glutamic acid (L, D-) cross-links, replacing the terminal D-alanine with the amino alcohol alaninol and cleaving γ-D-glutamyl-meso-diaminopimelic acid (D, D-) crosslinks, which reduce the release of immunogenic muropeptides, thereby limiting recognition by innate immune receptors, such as NOD1/2, and attenuating the inflammatory response [24], [25]. Additionally, in *Salmonella* Typhi, the elevated levels of L, D-crosslinks play a dual role by facilitating the secretion of the typhoid toxin [22]. UppP’s crucial function in peptidoglycan synthesis suggests that it may play a role in cell wall adaptation during stress, thereby enhancing *Salmonella*’s survival. In this study, we generated an *uppP* knockout strain of *S.* Typhimurium and evaluated its impact on bacterial survival and virulence. Our findings demonstrate that UppP is essential for *S.* Typhimurium proliferation in RAW 264.7 macrophages and systemic dissemination in C57BL/6 mice by promoting resistance to host-induced nitrosative stress.

## Results

### UppP affects bacterial cell dimensions and strength

To investigate the role of undecaprenyl pyrophosphate phosphatase in *Salmonella* pathogenesis, we generated an *uppP-*deleted strain of *Salmonella* Typhimurium by a one-step chromosomal gene inactivation method [26]. We observed that the *uppP* deleted strain demonstrated no significant difference in growth in nutrient-rich LB broth (Fig. 1A), M9 minimal media (Fig. 1B) and acidic F-media (pH-5.4, similar to the pH of the *Salmonella*-containing vacuole [SCV]) [27], [28] (Fig. 1C).

**Figure 1.**
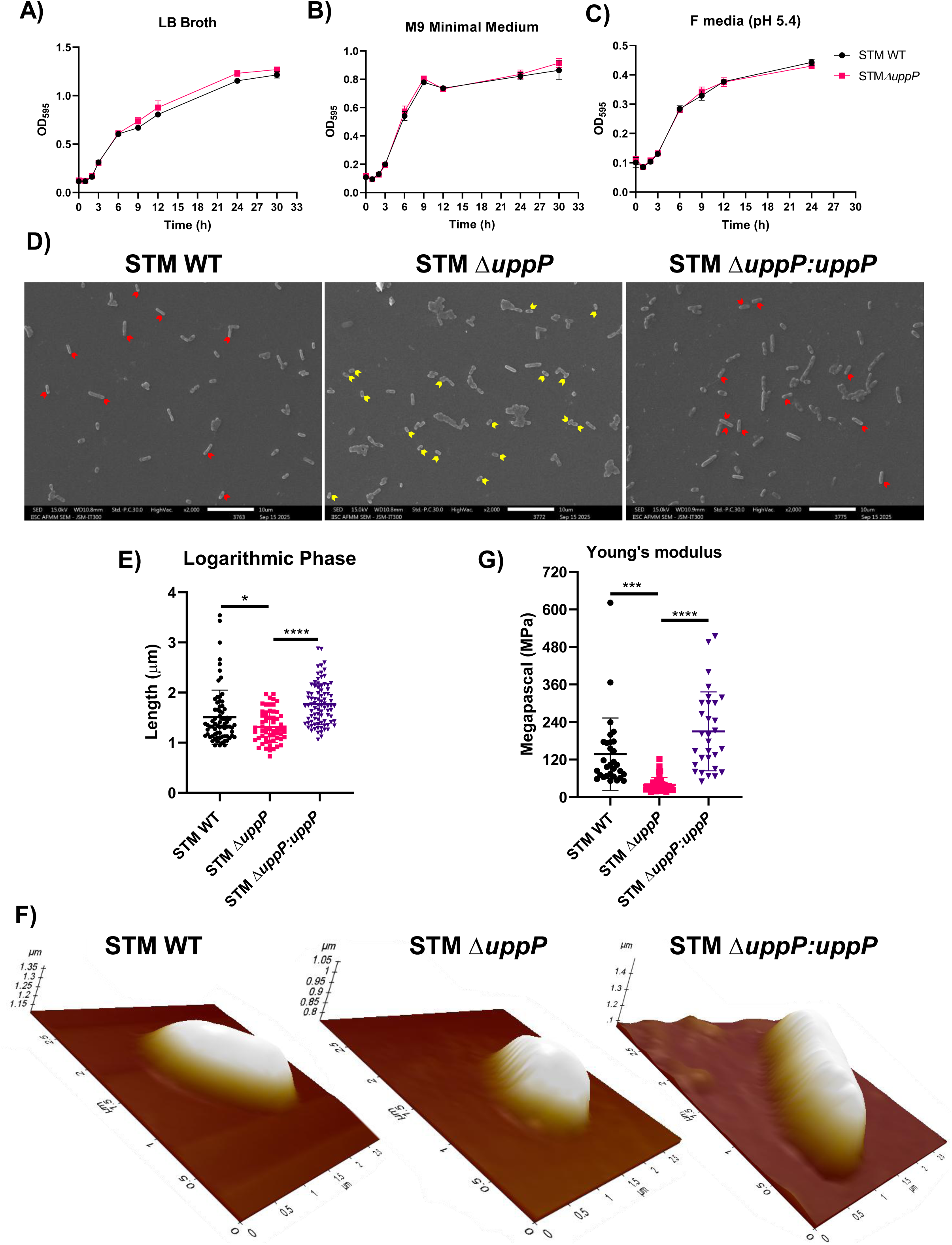
Loss of *uppP* affects the bacterial cell dimensions and strength A-C) Growth kinetics of STM WT and STM *ΔuppP* by measuring optical density at 595nm (OD595nm) in LB (Luria Bertani) broth (A), M9 minimal media (B), and acidic F-media (C). Data are representative of N=3, n=3 ± SD (standard deviation). D) Representative scanning electron micrographs of STM WT, STM *ΔuppP* and STM *ΔuppP*:*uppP* logarithmic phase cells (1:100 dilution). Red and yellow arrows represent cells with increased and decreased length, respectively. Scale bar = 5μm E) Quantification of the length of the logarithmic phase (1:100 dilution) bacterial cells captured in the SEM images (D). Data are representative of N=2, n≥50 ± SD (standard deviation). One-way ANOVA was performed to obtain the p-values. (****P ≤ 0.0001, ***P ≤ 0.001, **P ≤ 0.01, *P ≤ 0.05 ns- non-significant) F) Representative atomic force images of STM WT, STM *ΔuppP* and STM *ΔuppP*:*uppP*. G) Quantification of Young’s modulus (MPa- Megapascal) of the bacterial cells captured in the AFM images. Data are representative of n≥30 ± SD (standard deviation). One-way ANOVA was performed to obtain the p-values. (****P ≤ 0.0001, ***P ≤ 0.001, **P ≤ 0.01, *P ≤ 0.05 ns- non-significant)

A continuous supply of undecaprenyl phosphate is crucial for normal cell shape maintenance in *Escherichia coli*, and its depletion induces morphological defects [29], [30], [31]. To dissect the contribution of UppP in bacterial cell morphology of *Salmonella*, we performed Scanning Electron Microscopy (SEM) to examine the logarithmic (1:100 dilution) and stationary-phase cultures of STM WT and STM *ΔuppP*. The quantitative analysis of cell length revealed that STM *ΔuppP* (1.319 ± 0.3145 μm) exhibits reduced cell length in comparison to STM WT (1.509 ± 0.5406 μm) during the log phase, which was restored upon complementation of *uppP* (1.768 ± 0.4086 μm) (Fig. 1D-E). Whereas there is an increase in cell length of STM *ΔuppP* (1.236 ± 0.2295 μm) in comparison to STM WT (1.067 ± 0.2360 μm) during the stationary phase (S1A-B). Interestingly, when STM Δ*uppP* cells were subcultured at a higher dilution of 1:1000, allowing them to grow for a longer period in the logarithmic phase, we observed that the mutant (1.963 ± 0.5102 μm) became notably more elongated compared to the wild type (1.642 ± 0.4547 μm). Additionally, cell size returned to normal upon complementation (1.704 ± 0.4999 μm) (Fig. S1C-D).

Furthermore, we performed atomic force microscopy (AFM) to evaluate the stiffness of the bacterial cell envelope. Young’s modulus, quantified from AFM micrographs, was reduced in the STM Δ*uppP* (39.55 ± 23.26 MPa) strain compared to the wild type (137.5 ± 115.5 MPa), and this reduction was also restored upon complementation (210.3 ± 126.2 MPa) (Fig. 1F-G, S1E). Additionally, we evaluated the porosity of the outer membrane through a bisbenzimide uptake assay, as it is a major determinant of the stiffness of the bacterial cell envelope [32]. We observed no significant difference in fluorescence uptake between the wild-type and the mutant, either during the stationary or logarithmic phase (Fig. S1F). Our results indicate that *uppP* contributes to the maintenance of bacterial cell size and strength.

To investigate whether the observed defects in bacterial cell length and stiffness of STM Δ*uppP* can be attributed to biochemical changes in the cell envelope, we evaluated the composition of two major components: the peptidoglycan layer and lipopolysaccharide (LPS), both of which are dependent on undecaprenyl phosphate for their biosynthesis. The muropeptide composition of STM WT and STM Δ*uppP* analysed using HPLC revealed no significant differences in the muropeptide profile or peptidoglycan cross-links (Figure S2A-C). Similarly, silver staining of LPS showed no differences in the banding pattern or intensity between the wild-type and the mutant (Fig. S2D). Collectively, our results suggest that deletion of *uppP* has an impact on bacterial cell morphology and stiffness, without influencing the biochemical composition of either the peptidoglycan layer or lipopolysaccharide (LPS).

### *uppP* plays a role in the intracellular survival of *Salmonella* in RAW 264.7 macrophages

As macrophages play a crucial role in the dissemination of *Salmonella* within the host, we performed an intracellular survival assay using the murine macrophage-like cell line RAW 264.7. We assessed the percentage of phagocytosis/invasion, which represents the percentage of bacteria from the total pre-inoculum that invade the cell, and fold proliferation, which accounts for the extent of bacterial replication occurring inside the cell with respect to an early time point, respectively. We found that STM Δ*uppP* exhibits impaired invasion and fold proliferation compared to STM WT in RAW264.7 cells. The decreased percentage invasion and fold proliferation of the mutant is rescued upon complementation of the *uppP* gene (Fig. 2A, S3A). Since the intestinal epithelial barrier is *Salmonella*’s initial point of contact, we also evaluated the intracellular survival of STM Δ*uppP* in Caco-2 cells. However, we observed no significant difference in the invasion or fold proliferation of STM Δ*uppP* compared to wild-type in Caco-2 cells (Fig. S3B-C).

**Figure 2.**
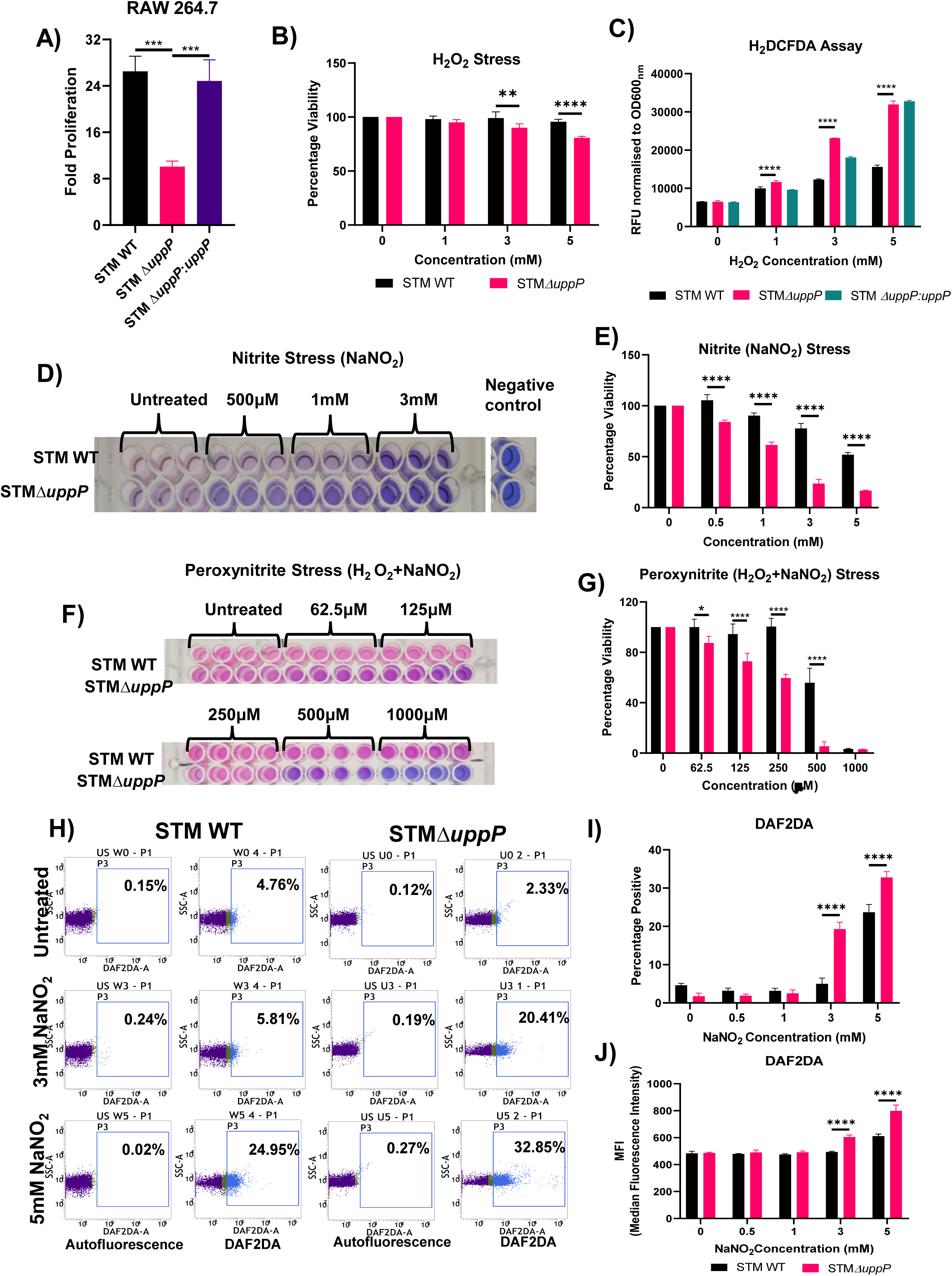
*uppP* facilitates *Salmonella* to withstand nitrite and peroxynitrite stress A) Fold proliferation of STM WT, STM *ΔuppP*, and STM *ΔuppP*:*uppP*, within RAW 264.7 macrophages (MOI=10). Results are mean of N=4, n=3 ± SEM (standard error of mean). One-way ANOVA was performed to obtain the p-values. (****P ≤ 0.0001, ***P ≤ 0.001, **P ≤ 0.01, *P ≤ 0.05 ns- non-significant) B) Percentage viability of STM WT and STM *ΔuppP* in the presence of varying concentrations of H2O2 measured by resazurin assay. Results are representative of N=2, n=4 ± SD. Two-way ANOVA was performed to obtain the p-values. (****P ≤ 0.0001, ***P ≤ 0.001, **P ≤ 0.01, *P ≤ 0.05 ns- non-significant) C) Estimation of reactive oxygen species (ROS) levels in STM WT, STM *ΔuppP* and STM *ΔuppP*:*uppP* by H2DCFDA staining in the presence of varying concentrations of H2O2. Results are representative of N=5, n=4 ± SD. Two-way ANOVA was performed to obtain the p-values. (****P ≤ 0.0001, ***P ≤ 0.001, **P ≤ 0.01, *P ≤ 0.05 ns- non-significant) D-E) Representative image (D) and percentage viability (E) of resazurin assay of STM WT and STM Δ*uppP* in the presence of varying concentrations of NaNO2 (*in vitro* nitrite stress). Results are representative of N=3, n=3 ± SD. Two-way ANOVA was performed to obtain the p-values. (****P ≤ 0.0001, ***P ≤ 0.001, **P ≤ 0.01, *P ≤ 0.05 ns- non-significant) F-G) Representative image (F) and percentage viability (G) of resazurin assay of STM WT and STM Δ*uppP* in the presence of varying concentrations of NaNO2 + H2O2 (*in vitro* peroxynitrite stress). Results are representative of N=2, n=4 ± SD. Two-way ANOVA was performed to obtain the p-values. (****P ≤ 0.0001, ***P ≤ 0.001, **P ≤ 0.01, *P ≤ 0.05 ns- non-significant) H-J) Estimation of nitric oxide in STM WT and STM Δ*uppP* by DAF2DA staining in the presence of varying concentrations of NaNO2 by flow cytometry. Representative SSCA vs DAF2DA dot plots of STM WT and STM Δ*uppP* (H). Quantification of DAF2DA positive percentage population (I) and median fluorescence intensity (MFI) (J). Results are representative of N=3, n=4 ± SD. Two-way ANOVA was performed to obtain the p-values. (****P ≤ 0.0001, ***P ≤ 0.001, **P ≤ 0.01, *P ≤ 0.05 ns- non-significant)

As SPI-1 can also contribute to macrophage invasion [33], we evaluated the transcript levels of SPI-1 genes. We observed increased *sipA* (SPI-1 effector), *invF*, *hilA* (transcriptional regulators), and *prgH* (structural translocon component) transcripts in the mutant during both the logarithmic (6h) and stationary phase (12h) (Fig. S3D-E). Increased expression of SPI-1 genes is also associated with macrophage apoptosis during *Salmonella* infection [34], [35], [36]. We hypothesised that the reduced invasion and fold proliferation of STM *ΔuppP* may be attributed to the infected cells undergoing apoptosis. To evaluate the same, 16 hours post-infection, RAW 264.7 cells were treated with tetracycline, followed by an MTT (3-(4,5-dimethylthiazol-2-yl)-2,5-diphenyltetrazolium bromide) assay to determine cell viability. No significant difference was observed in the cell viability between the wild-type, mutant and uninfected control (Fig. S3F). This suggests that apoptosis, or other forms of cell death, may not account for the decreased invasion and proliferation of STM Δ*uppP*.

*Salmonella* Pathogenicity Island (SPI)-2 is a key player in governing *Salmonella* survival in macrophages [37]. Since STM Δ*uppP* exhibited reduced fold proliferation in RAW 264.7 macrophages, we evaluated the transcript levels of SPI-2 effectors *sifA* and *spiC*. Interestingly, STM Δ*uppP* exhibited elevated transcripts of *sifA* and *spiC* during the stationary phase, compared to the wild-type (Fig. S3G).

Multiple host defences within the macrophage, such as acidic pH, reactive oxygen species, reactive nitrogen species, antimicrobial peptides, nutrient deprivation, etc, influence *Salmonella* survival [8]. We assessed the susceptibility of STM *ΔuppP* to exogenous stresses of hydrogen peroxide, nitrite and peroxynitrite by a resazurin dye-assisted cell viability assay. Despite exhibiting elevated reactive oxygen species (ROS) levels, as measured by H2DCFDA staining, STM Δ*uppP* displayed only a marginal reduction in viability when subjected to hydrogen peroxide stress (Fig. 2B-C). In contrast, treatment with increasing concentrations of sodium nitrite and peroxynitrite resulted in significantly decreased viability of STM Δ*uppP* compared to the wild-type. (Fig. 2D-G). Additionally, complementing *uppP* in the mutant strain restored bacterial viability when exposed to nitrite stress (Fig. S3H). We evaluated intracellular nitric oxide production in STM Δ*uppP* under sodium nitrite stress using DAF2DA staining. Flow cytometry-mediated visualisation showed that STM *ΔuppP* demonstrates an increased percentage of DAF2DA-positive cells upon nitrite stress (Fig. 2H-I). Moreover, on exposure to nitrite stress, STM *ΔuppP* exhibited an elevated median fluorescence intensity (MFI) compared to the wild type, indicating elevated nitric oxide levels (Fig. 2J). Our findings suggest that UppP contributes to *Salmonella* survival in RAW264.7 cells by aiding the bacterial defence against nitrosative stress.

### UppP deletion leads to increased nitric oxide (NO) production and nitrotyrosine localisation with intracellular bacteria within macrophages

To evaluate if the reduced fold proliferation of STM *ΔuppP* is attributed to nitrosative stress, we measured nitric oxide production by infected macrophages. We performed the Griess assay to evaluate the levels of extracellular NO in the culture supernatants of RAW 264.7 cells infected with STM WT, STM *ΔuppP* and STM *ΔuppP:uppP* for 16 hours. There was an 8 to 10-fold increase in the extracellular nitrite concentration in the culture supernatants of RAW 264.7 cells infected with STM *ΔuppP* (38.19 ± 5.025 μM) in comparison to the wild-type control (3.949 ± 1.418 μM). Moreover, upon complementation of *uppP*, the extracellular nitrite levels were restored (3.172 ± 1.006 μM), similar to that of the wild-type (Fig. 3A). We infected RAW 264.7 cells with STM WT and STM *ΔuppP,* transformed with the pFPV-mCherry plasmid, responsible for the constitutive expression of mCherry [38]. This was followed by DAF2DA staining 16h post-infection. The STM *ΔuppP*-mCherry infected cells showed increased MFI compared to STM WT-mCherry infected cells, indicating elevated nitric oxide levels (Fig. 3B-C, S4A-C). To verify that the elevated NO production was specifically STM *ΔuppP:uppP* for 16h, followed by DAF2DA staining. We observed increased intracellular NO levels in STM *ΔuppP-*infected cells, which were restored to wild-type levels upon complementation (Fig. S4D-E). NO exerts its bactericidal activity by oxidising to form NO adducts, such as nitrites and peroxynitrites, with peroxynitrite nitrosylating tyrosine and impairing protein function [39]. To assess this, we used confocal laser scanning microscopy to estimate nitrotyrosine levels in RAW 264.7 cells 16h post-infection with eGFP-tagged STM WT and STM *ΔuppP*. Our results show that in comparison to STM WT, there is an increase in the colocalisation of nitrotyrosine residues with STM *ΔuppP*. Moreover, cells infected with STM *ΔuppP* demonstrated increased levels of nitrotyrosine residues, consistent with our flow cytometry findings (Fig. 3D-F). To further solidify our findings, we performed an ELISA to quantify nitrotyrosine residues in the lysates of RAW 264.7 cells infected with STM WT, STM *ΔuppP* and STM *ΔuppP:uppP* for 16h. The cell lysates of STM *ΔuppP-*infected cells exhibited increased levels of nitrotyrosine residues, which reverted to wild-type levels upon complementation of *uppP* (Fig. S4F). Taken together, the elevated nitric oxide and nitrotyrosine levels in RAW 264.7 cells infected with STM *ΔuppP* suggest that *uppP* may have a protective role against nitrosative stress.

**Figure 3.**
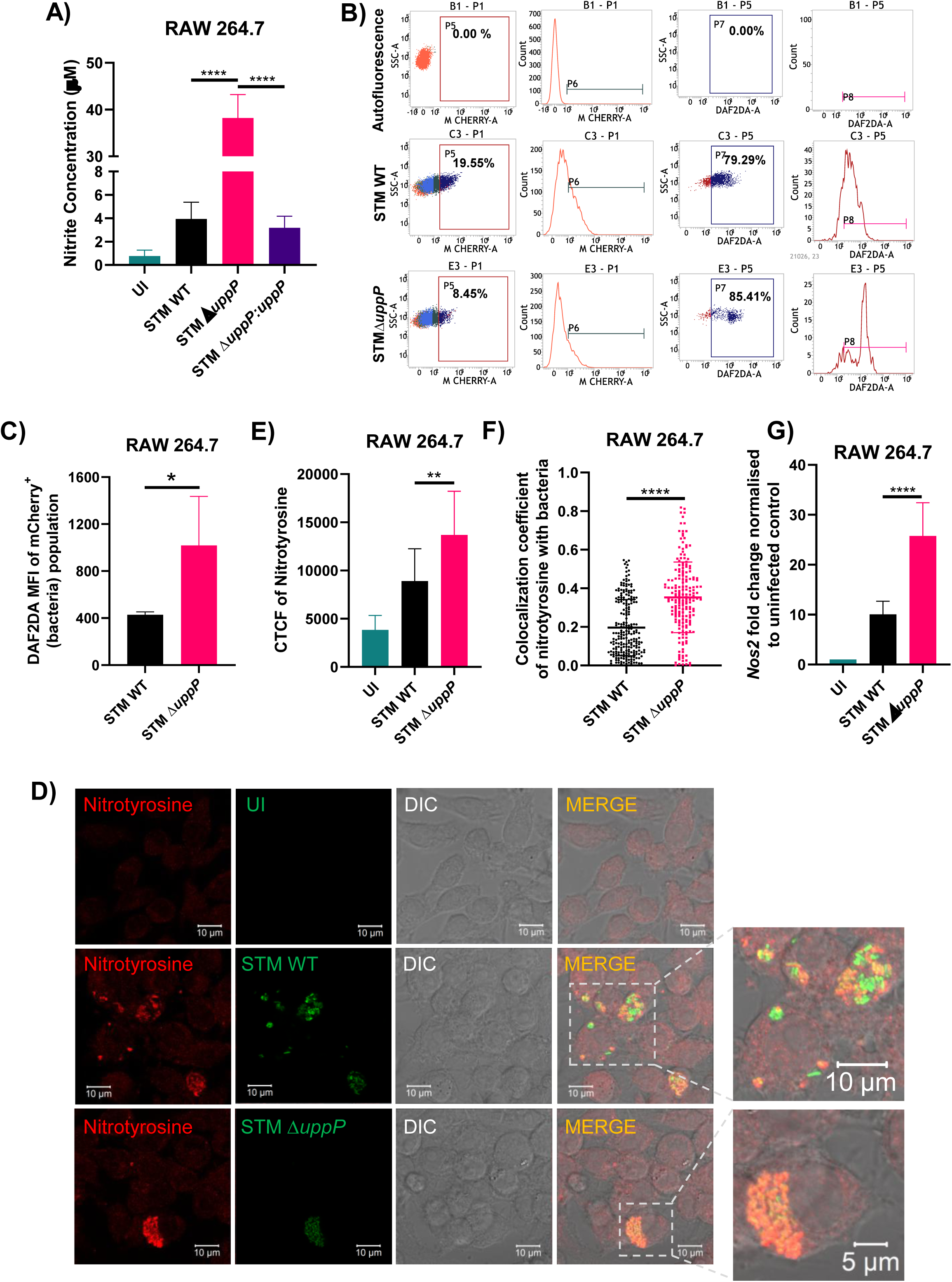
*uppP* sustains *Salmonella* survival in macrophages by countering host-induced nitrosative stress A) Estimation of extracellular nitrite by Griess assay in culture supernatants of RAW264.7 cells, 16h post-infection with STM WT, STM *ΔuppP* and STM *ΔuppP:uppP*, MOI=10, N=3, n=3 ± SD. One-way ANOVA was performed to obtain the p-values. (****P ≤ 0.0001, ***P ≤ 0.001, **P ≤ 0.01, *P ≤ 0.05 ns- non-significant) B-C) Estimation of nitric oxide in macrophages infected with mCherry-tagged STM WT and STM Δ*uppP* at MOI=20 by DAF2DA staining followed by flow cytometry. Representative mCherry vs SSC-A, DAF2DA vs SSC-A, dot plots and histograms of STM WT and STM Δ*uppP* (B). Quantification of DAF2DA median fluorescence intensity (MFI) of the mCherry-positive population (C). Results are representative of N=3, n=4 ± SD. An unpaired two-tailed Student’s t-test was performed to obtain the p-values. (****P ≤ 0.0001, ***P ≤ 0.001, **P ≤ 0.01, *P ≤ 0.05 ns- non-significant) D-F) Representative immunofluorescence images of RAW 264.7 cells infected with STM WT and STM *ΔuppP* for 16h, MOI=10, immunostained for nitrotyrosine (D). Quantification of corrected total cell fluorescence (CTCF) of nitrotyrosine in infected cells (E) and colocalisation of nitrotyrosine with bacteria (F). Results are representative of N=3, n≥50 ± SD. Unpaired two-tailed Student’s t-test was performed to obtain the p-values. (****P ≤ 0.0001, ***P ≤ 0.001, **P ≤ 0.01, *P ≤ 0.05 ns- non-significant) G) Quantification of *Nos2* mRNA expression in RAW264.7 cells infected with STM WT and STM Δ*uppP* for 16h. Results are representative of N=2, n=3 ± SD. One-way ANOVA was performed to obtain the p-values. (****P ≤ 0.0001, ***P ≤ 0.001, **P ≤ 0.01, *P ≤ 0.05 ns-non-significant)

Elevated NO production is also a prominent indicator of M1 polarised proinflammatory macrophages [40]. To determine whether the increased NO production resulted from a heightened immune response against the mutant, hence aiding in bacterial clearance, we quantified the levels of *Nos2* transcript and two other proinflammatory markers, CD86 and TNF-α, using flow cytometry and ELISA, respectively. The *Nos2* gene encodes for the enzyme inducible nitric oxide synthase (iNOS), which generates NO. We observed that the *Nos2* mRNA transcript was elevated in RAW 264.7 cells upon infection with STM *ΔuppP* in comparison to STM WT (Fig. 3G). Flow cytometry analysis showed that there was an increase in the MFI of STM *ΔuppP-*infected RAW 264.7 cells in comparison to the STM WT and STM *ΔuppP:uppP*-infected cells, indicating a higher cell surface expression of CD86 upon infection with STM *ΔuppP* (Fig. S5A-B). Additionally, we observed that the spent media of RAW 264.7 cells infected with STM Δ*uppP* showed a minimal increase in TNF-α levels compared to STM WT-infected cells (Fig. S5C). Altogether, these data indicate that STM Δ*uppP* may exhibit increased immunogenicity, inducing a stronger proinflammatory immune response within host macrophages in comparison to STM WT. This NO-driven immune response, along with the mutant’s innate sensitivity to nitrosative stress, may facilitate its clearance in macrophages.

### UppP mutant shows increased survival on alleviating nitrosative stress

To corroborate our findings of *uppP*’s role in withstanding nitrosative stress, we inhibited inducible nitric oxide synthase (iNOS) by an irreversible inhibitor, 1400W [27], [41], and observed a rescue in the fold proliferation of STM*ΔuppP* in RAW 264.7 cells (Fig. 4A).

**Figure 4.**
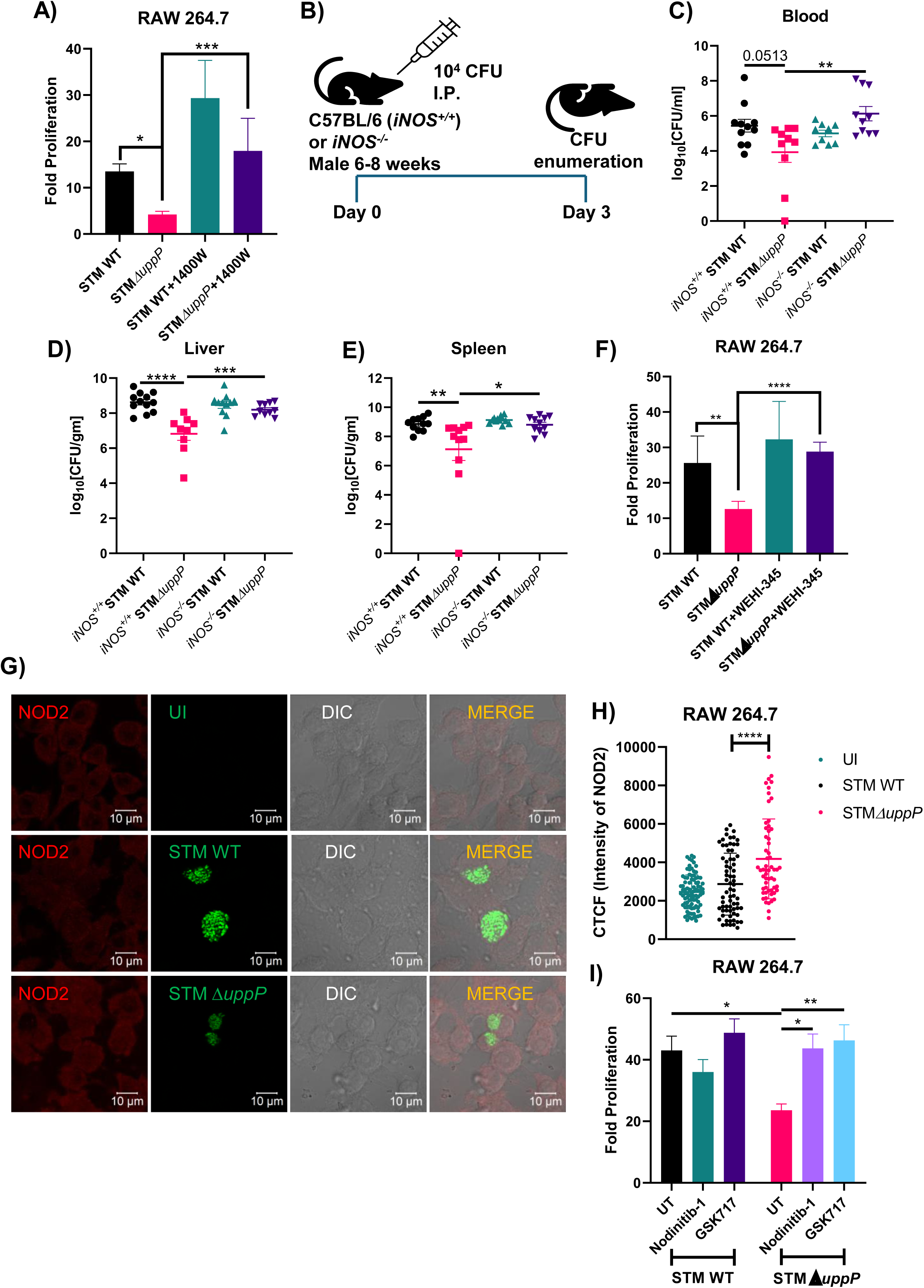
*uppP* counters nitrosative stress during systemic infection A) Fold proliferation of STM WT and STM *ΔuppP* within RAW 264.7 macrophages on treatment with 1400W MOI=10. Results are representative of N=3, n=3 ± SD (standard deviation). One-way ANOVA was performed to obtain the p-values. (****P ≤ 0.0001, ***P ≤ 0.001, **P ≤ 0.01, *P ≤ 0.05 ns- non-significant) B) Schematic representation of the experimental protocol followed for intraperitoneal (I.P.) and infection, enumerating bacterial burden in tissues C-E) Enumeration of bacterial burden in the blood (C), liver (D), and spleen (E) on infection via intraperitoneal injection in C57BL/6 (*iNOS^+/+^*) and *iNOS^-/-^* mice. Results are representative of N=2 n=7 ± SEM (standard deviation). Mann-Whitney test was performed to obtain the p values. (****P ≤ 0.0001, ***P ≤ 0.001, **P ≤ 0.01, *P ≤ 0.05 ns- non-significant) F) Fold proliferation of STM WT and STM *ΔuppP* within RAW 264.7 macrophages on treatment with WEHI-345 MOI=10. Results are representative of N=3, n=3 ± SD (standard deviation). One-way ANOVA was performed to obtain the p-values. (****P ≤ 0.0001, ***P ≤ 0.001, **P ≤ 0.01, *P ≤ 0.05 ns- non-significant) G-H) Representative immunofluorescence images of RAW 264.7 cells infected with STM WT and STM *ΔuppP* for 16h, MOI=10, immunostained for NOD2 (G). Quantification of corrected total cell fluorescence (CTCF) of NOD2 in infected cells (H). Results are representative of N=2, n≥50 ± SD. One-way ANOVA was performed to obtain the p-values. (****P ≤ 0.0001, ***P ≤ 0.001, **P ≤ 0.01, *P ≤ 0.05 ns- non-significant) I) Fold proliferation of STM WT and STM *ΔuppP* within RAW 264.7 macrophages on treatment with Nodinitib-1 and GSK717, MOI=10. Results are mean of N=3, n=3 ± SEM (standard error of mean). One-way ANOVA was performed to obtain the p-values. (****P ≤ 0.0001, ***P ≤ 0.001, **P ≤ 0.01, *P ≤ 0.05 ns- non-significant)

To investigate the role of *uppP* in *Salmonella* pathogenesis *in vivo*, C57BL/6 mice were orally challenged with a dose of 10^7^ CFU of STM WT, and STM *ΔuppP* and bacterial burden was assessed 5 days post-infection (Fig. S6A). STM *ΔuppP* displayed reduced dissemination by blood and decreased bacterial loads in the intestine, MLN (mesenteric lymph node), liver and spleen (Fig. S6B-F). Additionally, STM *ΔuppP*-infected mice demonstrated a modest increase in survival compared to STM WT (Fig. S6G). Furthermore, to evaluate whether the mutant’s attenuation in the mouse model was due to its increased susceptibility to nitrosative stress, *iNOS^-/-^*C57BL/6 mice were orally challenged with 10^5^ CFU of STM WT and STM *ΔuppP* (Fig. S6H). Notably, STM *ΔuppP* still exhibited attenuation in dissemination and reduced organ burden compared to the wild type (Fig. S6I-M). These findings suggest that UppP may have additional roles in bacterial invasion of the intestine, beyond its role in conferring resistance to nitrosative stress. To specifically evaluate UppP role in countering nitrosative stress during systemic infection, *iNOS^+/+^* and *iNOS^-/-^* C57BL/6 mice were intraperitoneally injected with 10^4^ CFU of STM WT and STM *ΔuppP* (Fig. 4B). We observed a significantly increased burden of STM *ΔuppP* in the blood, liver and spleen of *iNOS^-/-^* mice compared to *iNOS^+/+^* mice, substantiating the role of UppP in shielding *Salmonella* from nitrosative stress (Fig. 4C-E).

Induction of iNOS and NO production is triggered by downstream signalling from multiple pattern recognition receptors (PRRs). These include but are not limited to TLR4, TLR2, NOD1, and NOD2, out of which the latter of the three receptors detects various components associated with the bacterial peptidoglycan layer and TLR4 recognises lipopolysaccharide (LPS) [42], [43], [44], [45]. Also, both NOD1 and NOD2 have been reported to localise on *Salmonella-*containing vacuole [46], [47]. As UppP is involved in the biosynthesis of both peptidoglycan and LPS, we aimed to determine whether signalling through their cognate PRRs contributes to the enhanced susceptibility of STM *ΔuppP* in macrophages. We inhibited TLR2, TLR4 and RIPK2 (the immediate downstream kinase of NOD receptors) using small molecule inhibitors C29, Restoravid and WEHI-345, respectively [48], [49], [50]. We observed a rescue in the fold proliferation of STM *ΔuppP* in RAW 264.7 cells upon treatment with RIPK2 inhibitor WEHI-345, but not with either C29 or Restoravid, suggesting that NOD signalling may play a role in the increased susceptibility of STM *ΔuppP* in macrophages (Fig. 4F, S7A). Furthermore, to identify the NOD receptor which may be involved, we estimated the transcript levels of *Nod1* and *Nod2* in RAW264.7 cells infected with STM WT and STM *ΔuppP*. Although we did not observe any significant difference in the transcript levels of *Nod1*, we detected elevated *Nod2* transcript levels in RAW 264.7 cells, 2 hours post-infection with STM *ΔuppP* (Fig. S7B-C).

We performed immunofluorescence microscopy in STM WT and STM *ΔuppP-*infected RAW264.7 cells to extend our findings to the translational level. We found that STM *ΔuppP-*infected RAW264.7 cells at 6 and 16 hours post-infection exhibited a higher corrected total cell fluorescence (CTCF) of NOD1 and NOD2, respectively, compared to the wild-type and uninfected control (Fig. 4G-H, S7D-E). To further validate our findings, we treated RAW 264.7 cells with Nodinitib-1 and GSK717, which inhibit NOD1 and NOD2, respectively [51]. We observed a rescue in the fold proliferation of STM *ΔuppP* in RAW264.7 cells, treated with either Nodinitib-1 or GSK717, suggesting that both NOD1 and NOD2-mediated signalling may contribute to the attenuated proliferation of STM *ΔuppP* within macrophages (Fig. 4I).

## Discussion

Cell wall integrity is vital for bacterial structure, protection, and survival. Reports in *Escherichia coli*, *Salmonella* Typhimurium and *Bacillus subtilis* have shown that cell size and shape are affected by changes in growth rate and nutritional availability [52], [53], [54], [55]. Although the STM Δ*uppP* mutant exhibited similar growth rate kinetics to the wild type in LB medium, similar to the observations of *uppP* mutant in *E*. *coli* [15]. However, we found structural defects in STM Δ*uppP* through SEM and AFM-assisted visualisation. Quantitative analysis of SEM images confirmed a biphasic effect: STM *ΔuppP* cells were shorter in logarithmic phase (1:100 dilution) but longer in stationary phase than wild-type cells.

Interestingly, STM Δ*uppP* cells were notably more elongated than wild-type cells when allowed to grow in a longer logarithmic phase (1:1000 dilution). This may stem from altered surface area to volume (SA/V) ratios; increased SA/V in the low dilution (1:100) log phase could permit division at smaller sizes, while the reduced surface area synthesis rate in the stationary phase and the high dilution (1:1000) logarithmic phase might facilitate cell lengthening, resulting in a decrease in the SA/V ratio [56], [57]. Additionally, the AFM results revealed a reduced Young’s modulus of STM Δ*uppP* cells. Undecaprenyl phosphate, produced by the undecaprenyl pyrophosphate phosphatase activity of *uppP*, is a shared lipid carrier in the biosynthesis pathways of peptidoglycan, O antigen and enterobacterial common antigen (ECA), all of which contribute to the stiffness of the cell envelope to varying degrees [30], [31]. Recent work has demonstrated that the outer membrane comprising O antigen and ECA is a major determinant of mechanical stiffness in Gram-negative bacteria [32]. In conjunction with our SEM and AFM findings, this led us to hypothesise that STM Δ*uppP* may have an altered cell envelope. Interestingly, the biochemical analysis of the composition of both the peptidoglycan and LPS revealed no significant differences between the mutant and the wild-type strains. This suggests that the reduced Young’s modulus of the mutant and its biphasic morphological defects may be attributed to the availability dynamics of free undecaprenyl phosphate, which may in turn affect the biosynthesis rate of these macromolecules, rather than their composition or structure. Moreover, our bisbenzimide uptake assay demonstrated that there was no increased outer membrane porosity in the mutant, suggesting that the loss of *uppP* impairs the mechanical properties of the cell envelope without affecting the outer membrane’s barrier function.

The changes observed in the bacterial cell dimensions of STM Δ*uppP* may also contribute to its attenuated virulence. Maintaining the correct cellular shape is imperative for the pathogenicity of various Gram-negative bacteria. For example, peptidoglycan hydrolase deletion mutants of *Helicobacter pylori* and *Campylobacter jejuni* deviate from their standard helical shape to a rod or curved rod morphology, which leads to impaired colonisation in the mouse and chick model, respectively [58], [59], [60]. Furthermore, Uropathogenic *E. coli* (UPEC) subverts neutrophil-mediated phagocytosis by undergoing filamentation [61]. Thus, the morphological alterations in STM Δ*uppP* are likely to influence its survival in *in vitro* and *in vivo* settings.

To further explore the functional consequence of UppP during *Salmonella* pathogenesis, we infected RAW 264.7 murine macrophages and Caco-2 colon carcinoma cells with the STM Δ*uppP*. Though STM Δ*uppP* demonstrated no growth defects in Caco-2, its proliferation was attenuated in RAW 264.7 cells. Resazurin-assisted live/ dead assay revealed that STM Δ*uppP* had increased susceptibility to nitrosative stress. Moreover, inhibition of inducible nitric oxide synthetase restored the proliferation of STM Δ*uppP* in macrophages. Even though

*Salmonella* avoids trafficking of iNOS-containing vesicles to the *Salmonella*-containing vacuole (SCV) in an SPI-2-dependent manner, the freely diffusible nature of nitric oxide (NO) can facilitate bacterium killing [62], [63]. The altered cell structure of the null mutant may further facilitate the entry of NO within the bacterium. Also, STM Δ*uppP-*infected macrophages exhibited elevated levels of NO and CD86, along with increased secretion of TNF-α, suggesting that the mutant elicits a robust proinflammatory immune response within macrophages. This enhanced immune response, driven by increased NO production and the mutant’s inherent nitrite sensitivity, may promote its attenuated survival within macrophages.

The nucleotide oligomerisation domain (NOD) - like receptors 1 and 2 (NOD1/2) are cytoplasmic pattern recognition receptors (PRRs) recognising γ-d-glutamyl-meso-diaminopimelic acid (iE-DAP) and muramyl dipeptide (MDP), which are constituents of the peptidoglycan layer [64]. NOD1 and NOD2, and RIPK2 (downstream effector protein kinase) are reported to colocalise with the SCV, along with endosomal transporters SLC15A3 and SLC15A4, which translocate MDP to NOD receptors [46], [47]. In the *Salmonella* colitis model, mice deficient for NOD1 and NOD2 have higher bacterial loads in the mucosal tissue [65]. *Mycobacterium tuberculosis* growth in human macrophages is controlled by NOD2-dependent NO production [66]. Similarly, NOD2 induces NO, which mediates bacterial clearance in the mouse model of *Chlamydophila pneumoniae-*induced pneumonia [67].

NOD1-induced NO production provides resistance against *Trypanosoma cruzi* in the murine model [68]. TLR2 and TLR4, which recognise peptidoglycan and LPS, respectively, also signal the production of NO during bacterial infections [42], [45]. Upon inhibiting RIPK2, NOD1 or NOD2, but not TLR2 and TLR4, we observed a rescue in the proliferation of STM Δ*uppP* in macrophages. Undecaprenyl phosphate is a common substrate for synthesising peptidoglycan, O antigen, and ECA, all of which function as essential virulence determinants [17]. Initially, we hypothesised that an insufficient influx of undecaprenyl phosphate may lead to the accumulation of precursors and intermediates, which may act as ligands to directly or indirectly activate PRRs. In our study, upon deletion of *uppP,* we did not observe any difference in the composition of either LPS or peptidoglycan. We propose an alternative hypothesis that the deletion of *uppP* may slow the synthesis rate of both peptidoglycan and lipopolysaccharide; however, the synthesis of the latter will be more significantly impacted than that of the former, as peptidoglycan is more essential to the cell envelope. The slowed synthesis of peptidoglycan and LPS may explain the mutant’s reduced stiffness and altered morphology, leading to its lysis more quickly within macrophages. Furthermore, upon bacterial lysis of the *uppP* mutant, the peptidoglycan fragments may be relatively more abundant, leading to the preferential activation of NOD receptors and driving macrophage activation and elevated NO levels.

In C57BL/6 mice with functional iNOS (*iNOS^+/+^*), intraperitoneal infection with STM Δ*uppP* showed reduced dissemination by blood and decreased colonisation in the spleen and liver.

This phenotype was reversed in iNOS-deficient mice, confirming the role of UppP in promoting *Salmonella* dissemination and colonisation by counteracting nitrosative stress.

To the best of our knowledge, our study is one of the first to report a unique function of undecaprenyl pyrophosphate phosphatase in mitigating nitrosative stress in *Salmonella* (Fig. 5), in addition to its canonical roles in cell envelope biogenesis and maintenance of morphology. Our findings potentially highlight undecaprenyl phosphate metabolism as a promising target for developing new antimicrobial strategies to enhance the efficacy of macrophage-mediated clearance of *Salmonella* infection.

**Figure 5.**
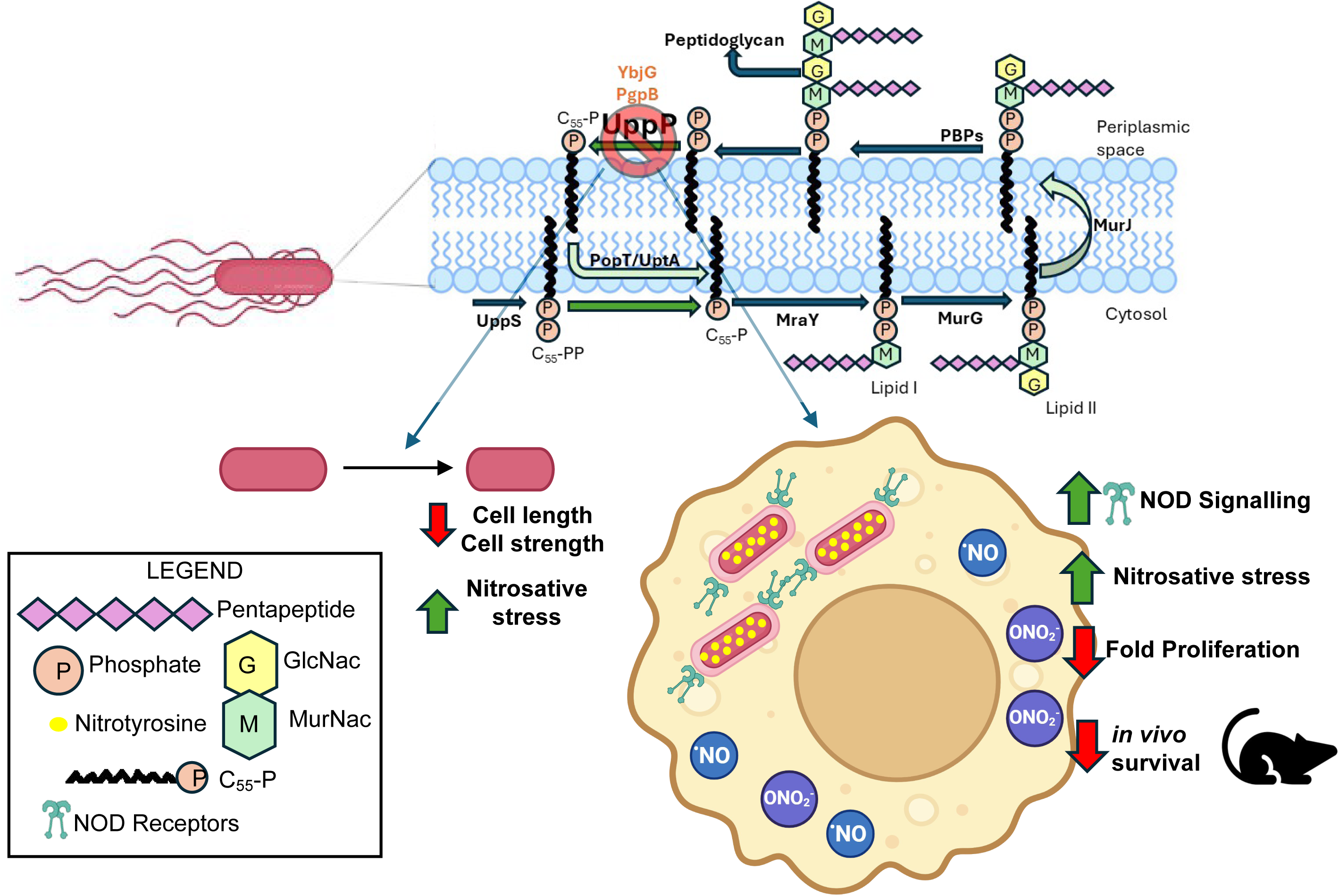
Graphical Abstract Increased susceptibility of STM *ΔuppP* to host nitrosative stress is indicated by elevated NO levels in RAW 264.7 cells and a greater accumulation of nitrotyrosine residues on the mutant. The elevated NO levels may be due to activation of NOD receptors, which is one of the factors contributing to STM *ΔuppP*’s reduced fold proliferation in RAW 264.7 cells and attenuated survival in the mouse model. NO: Nitric oxide, ONO2^-^: Peroxynitrite, NOD: Nucleotide-binding oligomerisation domain-containing protein, GlcNac: N-acetylglucosamine, MurNac: N-Acetylmuramic acid. Created with BioRender.com

## Material and methods

### Bacterial strains and growth condition

*Salmonella enterica* serovar Typhimurium 14028S (STM WT) was used in all experiments. Bacterial strains were cultured in Luria Bertani (LB) broth (HiMedia) with constant shaking (160 rpm) at 37°C (wild type, knockout strains) or 30°C (strains having temperature-sensitive pKD46 plasmid). Ampicillin (50μg/ml), Kanamycin (50μg/ml), Chloramphenicol (25μg/ml) and IPTG (500μM) were used wherever required. A complete list of bacterial strains and plasmids used in this study is provided in Table 1.

**Table 1.** Strains and plasmids used in this study

| Strains/ Plasmids | Characteristics | Source/ Information |
| --- | --- | --- |
| *Salmonella enterica* subspecies *enterica* serovar Typhimurium strain 14028S | Wild type (WT) | Gift from Prof. M. Hensel (Division of Microbiology, University of Osnabrück, Germany) |
| STMΔ*uppP* | Chl<sup>R</sup> | This study |
| STM Δ*uppP::uppP* (STMΔ*uppP::pQE60::uppP*) | Chl<sup>R</sup>, Amp<sup>R</sup> | This study |
| STM:pFPV-mCherry | Constitutive expression of mCherry in STM WT, Amp<sup>R</sup> | Laboratory stock |
| STMΔ*uppP::pFPV-mCherry* | Constitutive expression of mCherry in STM Δ*uppP*, Amp<sup>R</sup>, Chl<sup>R</sup> | This study |
| STM:pFPV25.1 | Constitutive expression of GFP in STM WT, Amp<sup>R</sup> | Laboratory stock |
| STMΔ*uppP::pFPV25.1* | Constitutive expression of GFP in STM Δ*uppP*, Amp<sup>R</sup>, Chl<sup>R</sup> | This study |
| pKD3 | Plasmid with FRT flanked by Chloramphenicol-resistance gene, Chl<sup>R</sup> | Laboratory stock |
| pKD46 | Plasmid expressing λ red recombinase system, Amp<sup>R</sup> | Laboratory stock |
| pQE60 | Low copy number plasmid, Amp<sup>R</sup> | Laboratory stock |
| pFPV-mCherry | Constitutive expression of mCherry, Amp<sup>R</sup> | Laboratory stock |
| pFPV25.1 | Constitutive expression of GFP, Amp<sup>R</sup> | Laboratory stock |

### Construction of *uppP* knockout strain of *Salmonella*

The knockout strain of *Salmonella* enterica serovar Typhimurium (strain 14028S) was generated using the one-step chromosomal gene inactivation method developed by Datsenko and Wanner [26]. In a concise overview, STM WT was transformed with pKD46 plasmid, housing a ’lambda red recombinase system’ controlled by an arabinose-inducible promoter. The transformed cells were cultivated in LB broth with ampicillin (50 μg/mL) and 50 mM arabinose at 30°C until an optical density of 0.35 to 0.4 at 600nm. Electrocompetent STM pKD46 cells were prepared by washing the bacterial cell pellet thrice with double autoclaved chilled Milli Q water and 10% (v/v) glycerol. Finally, the electrocompetent STM pKD46 cells were resuspended in 50 μL of 10% glycerol. For the knockout procedure, the chloramphenicol-resistant gene cassette (Chl^R^, 1.1 kb, used for knocking out *uppP*) was amplified from the pKD3 plasmid, utilising knockout primers. The amplified Chl^R^ gene cassettes underwent chloroform-isopropanol extraction and were electroporated into STM WT pKD46. The transformed cells were then plated on LB agar with chloramphenicol (25 μg/mL) to select the STM*ΔuppP* strain. Subsequently, the plates were incubated overnight at 37°C. The confirmation of knockout colonies was achieved through expression primers. The primers used for generating the knockout strain are listed in Table 2.

**Table 2.**
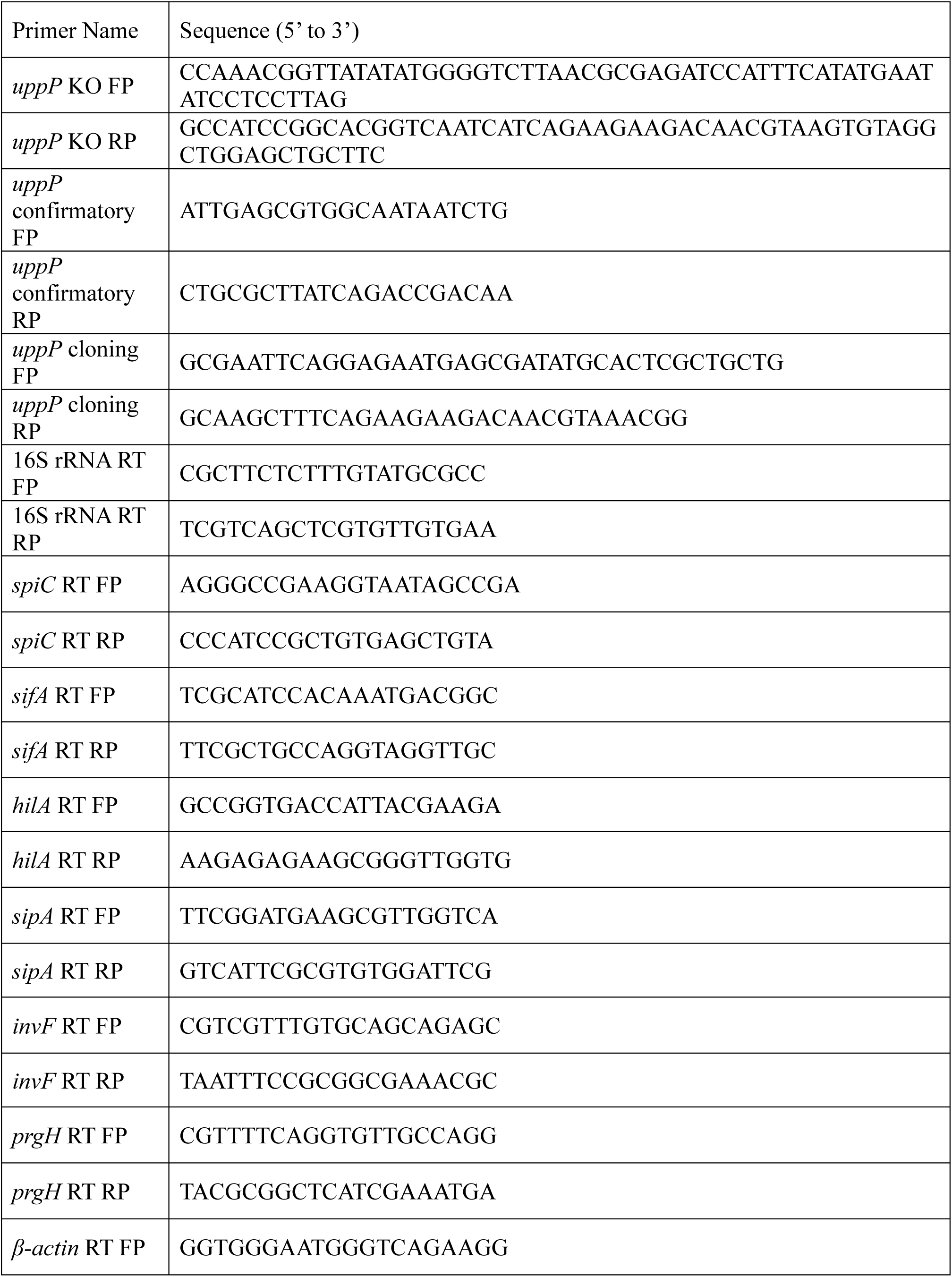
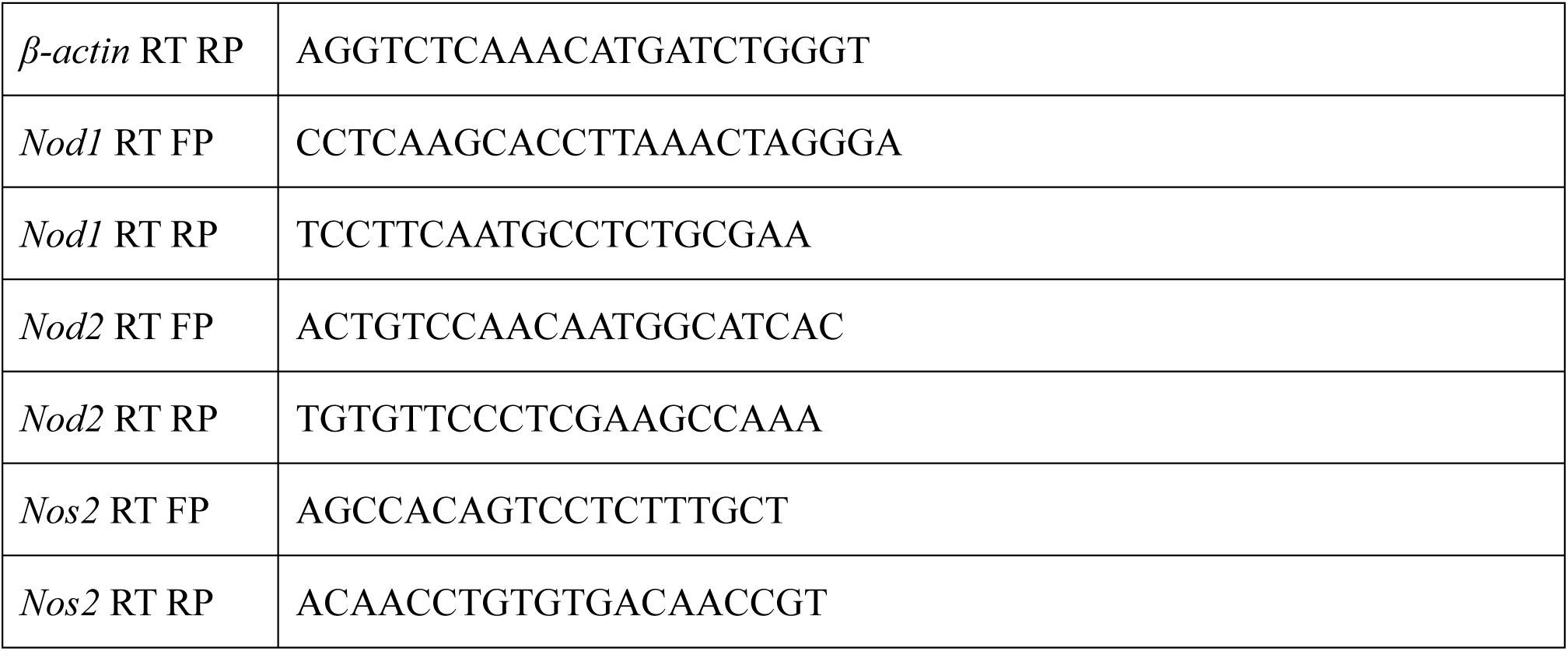
List of primer sequences (5’ to 3’)

### Construction of *uppP* complemented strain of *Salmonella*

The *uppP* gene was amplified using cloning primers through colony PCR. The resulting PCR product, purified via chloroform extraction, was then subjected to EcoRI (NEB) and HindIII (NEB) restriction digestion in CutSmart buffer (NEB) at 37°C overnight, alongside the empty pQE60 vector backbone. The double-digested insert and vector were ligated using T4 DNA ligase in 10X ligation buffer (NEB) overnight at 16°C. The ligated product was subsequently transformed into the knockout strains to generate the complemented strain. Verification of complementation was accomplished through the restriction digestion of the recombinant plasmid and Sanger sequencing. The primers used for generating the complement strain are listed in Table 2.

### Growth Curve Analysis

For *in vitro* growth curve experiments, a single colony of each strain was inoculated in Luria Bertani (LB) broth (HiMedia) with constant shaking (160 rpm) at 37°C, with the corresponding antibiotic. Overnight-grown stationary phase bacteria were sub-cultured at a ratio of 1: 100 in freshly prepared LB broth, M9 minimal media and F-media (50mM KCl, 75mM (NH4)SO4, 5mM K2SO4 10mM K2HPO4, 1Mm Bis-Tris, 8μM MgCl2, 38mM glycerol and 0.1% cas amino acids, pH- 5.4) kept at 37°C, with constant shaking (160 rpm). At different time intervals, aliquots were taken to measure [OD]595nm by TECAN 96 well microplate reader or for CFU analysis.

### Scanning electron microscopy (SEM)

Overnight cultures were subcultured in LB broth at a 1:100 or 1:1000 ratio for 5-6 hours to obtain logarithmic-phase cultures. Logarithmic phase bacterial cultures were drop-casted and immobilised over coverslips by air-drying, followed by overnight fixation with 3.5% glutaraldehyde at 4°C. The following day, the samples were washed with MilliQ water and treated with an increasing ethanol gradient (10%, 30%, 50%, 70%, 90%, and 100%) to facilitate dehydration. The samples were desiccated for 30 minutes, mounted on copper stubs using carbon tape and coated with gold particles under high vacuum.

Images were acquired with an ESEM Quanta microscope at a magnification of 5000X. ImageJ was used to analyse bacterial size.

### AFM (Atomic force microscopy)

AFM-mediated assessment of bacterial morphology was performed by adopting the protocol from [69]. Overnight cultures were subcultured in LB broth at a 1:100 ratio for 5-6 hours to obtain logarithmic-phase cultures. Logarithmic phase bacterial cultures were drop-casted and immobilised over coverslips by air-drying, followed by fixation with 3.5% paraformaldehyde. The samples were washed thrice with MilliQ water, followed by desiccation for 30 minutes. Images were acquired from an NX10 AFM. XEI software was utilised to calculate Young’s modulus.

### Bisbenzimide assay to determine the outer membrane porosity of bacteria

The outer membrane porosity of the bacterial strains (STM WT, STM Δ*uppP*, and STM Δ*uppP:uppP*) in both the stationary phase and logarithmic phase was measured using bisbenzimide (Sigma-Aldrich), as modified from a previously specified protocol [27], [70]. At the end of the incubation period, the OD600nm of the cultures was adjusted to 0.1 with sterile 1x PBS. Subsequently, 180 μL of 0.1 OD culture was added to a 96-well microplate containing 20 μl of bisbenzimide (10 μg/mL) solution and further incubated for 10 minutes at 37°C in a shaker incubator. Due to enhanced outer membrane porosity, when bisbenzimide is taken up by bacterial cells, it binds to the bacterial DNA and begins to fluoresce. The fluorescence intensity of DNA-bound bisbenzimide was measured in a TECAN 96-well microplate reader using 346 nm excitation and 460 nm emission filter.

### Peptidoglycan isolation and analysis

The isolation and analysis of peptidoglycan were performed as described in [71], [72], [73]. Overnight bacterial cultures were subcultured at a 1:100 dilution in 500 mL of LB broth. Bacterial strains were grown till the logarithmic phase (OD600nm ≈ 1) and subsequently harvested by centrifugation at 8000 RPM for 20 minutes at 4°C. The cell pellet was resuspended in chilled MQ water, and the suspension was added dropwise to an equal volume of a constantly boiling solution of 8% SDS. The suspension was boiled for an additional 40 minutes, with continuous stirring and allowed to cool overnight at RT. The sample was mixed with MQ water and pelleted by ultracentrifugation at 3,00,000g, for 90 minutes, at RT. After multiple washes to completely remove SDS, the pellet was resuspended in 10 mM Tris Cl pH 7, 10mM NaCl buffer and treated with 0.1 mg/mL of α-amylase for 2 hours at 37°C, followed by treatment with 0.2 mg/mL of pronase for 2 hours at 60°C. The enzymes were inactivated by boiling with an equal volume of 8% SDS for 20 minutes, followed by removal of SDS, with several washes of MQ water by centrifugation at 25,000g at RT. The peptidoglycan pellets were finally resuspended in 25 mM Tris-HCl, pH 8, and stored at -30°C.

For analysing the peptidoglycan composition, muropeptide fragments were obtained by digesting the peptidoglycan with 10 U of mutanolysin in a 25 mM Tris-HCl buffer, pH 8, at 37°C for 16 hours. The samples were centrifuged at 25,000g for 10 minutes at RT, and the supernatant comprising the soluble muropeptides was reduced with 1 mg of sodium borohydride in 50 mM sodium-borate buffer, pH 9, for 30 min. 88% ortho-phosphoric acid was added to decrease the pH to 2, thereby eliminating the excess borohydride. The sample was centrifuged to remove any additional particulate matter, and the supernatant was injected into a preheated (55°C) Zorbax 300SB RP-C18 column, connected to an Agilent Technologies RRLC 1200 HPLC system. Muropeptides were eluted using a 0 to 20% gradient of acetonitrile containing 0.1% trifluoroacetic acid with a flow rate of 0.5 mL/min for 80 minutes. Muropeptides were detected by measuring absorbance at 205 nm.

### LPS isolation and quantification

The isolation and quantification of LPS were performed according to the protocol described by Mahalakshmi et al [74]. Cell lysates were prepared by sonication of logarithmic-phase strains (STM WT, STM Δ*uppP*, and STM Δ*uppP:uppP*) resuspended in 10mM HEPES buffer (pH 7.4). The protein concentration of the cell lysates was estimated using the Bradford Reagent (Bio-Rad) with standard solutions comprising bovine serum albumin (BSA) concentrations ranging from 50 μg/μL to 0.31 μg/μL. Equal volumes of cell lysates, normalised to protein content, were mixed with an equal volume of tricine sample buffer (comprising 100 mM Tris-HCl, pH 6.8, 24% w/v glycerol, 8% w/v SDS, 5% v/v 2-β-mercaptoethanol, and 0.02% w/v bromophenol blue) and boiled for 10 minutes. 10 μL of proteinase K solution (2.5 mg/mL in sample buffer) was added to 50 μL of the boiled sample, followed by incubation at 60°C for 1 hour. Subsequently, the sample was centrifugation at 16,000 g for 30 minutes. The supernatants were loaded onto 16.5% tricine-SDS polyacrylamide gels, and LPS was visualised through silver staining.

### Cell culture protocol

DMEM- Dulbecco’s Modified Eagle Medium (Lonza) supplemented with 10% FBS (Gibco) was used to culture RAW 264.7 murine macrophages and the human colorectal adenocarcinoma cell line, Caco-2. The cells were maintained at 37°C in a humified incubator with 5% CO2. Additionally, Caco-2 cells were supplemented with 1% sodium pyruvate (Sigma) and 1% non-essential amino acids (Sigma). Cells were seeded at a density of 1.5 x 10^5^ cells per well in a 24-well plate for intracellular survival assay preceding infection.

### Gentamicin protection assay

Macrophages or epithelial cells were infected with stationary-phase or logarithmic phase bacterial culture with an MOI of 10. For synchronisation of the infection and enhancing bacterial adhesion, tissue culture plates were centrifugated at 600 rpm for 5 minutes, followed by incubation for 25 minutes at 37°C in a humified incubator with 5% CO2. Post-incubation cells were washed with 1X PBS, and fresh media (DMEM + 10% FBS) containing 100μg/ml gentamicin was added, followed by 1h of incubation at 37°C and 5% CO2. Subsequently, the gentamicin concentration was reduced to 25μg/ml and maintained until the cells were harvested.

### Percentage phagocytosis and Intracellular proliferation

Following infection of cells with STM at an MOI of 10, cells were treated with DMEM (Sigma) supplemented with 10% FBS (Gibco) containing 100μg/ml gentamicin for 1h. Subsequently, the gentamicin concentration was reduced to 25μg/ml and maintained until the specified time point.

Additionally, in some experiments, 1400W dihydrochloride (Sigma), an inhibitor for inducible nitric oxide synthetase (iNOS), or WEHI-345 (MedChemExpress, HY-18937), a RIPK2 inhibitor which delays NOD signalling [50], Nodinitib-1 (MedChemExpress, HY-18639), an inhibitor of NOD1, and GSK717 (MedChemExpress, HY-136555), an inhibitor of NOD2, were administered to cells at concentrations of 10μM, 1μM, 5μM, and 5μM respectively, alongside 25μg/ml gentamicin treatment. TLR2 and TLR4 were inhibited using C29 (MedChemExpress, HY-100461) and Resatorvid (MedChemExpress, HY-11109), respectively, at concentrations of 25 μM and 100 nm. The cells were treated with either C29 or Resatorvid for 4 hours prior to infection, followed by the addition of the inhibitor alongside the 25μg/ml gentamicin during the entire infection period. 2h and 16h post-infection; cells were lysed in 0.1% triton-X-100. Lysed cells and the pre-inoculum were serially diluted and plated onto LB agar or *Salmonella-Shigella* agar to obtain colony-forming units (CFU). Calculations for percentage phagocytosis/ invasion and fold proliferation are as follows:

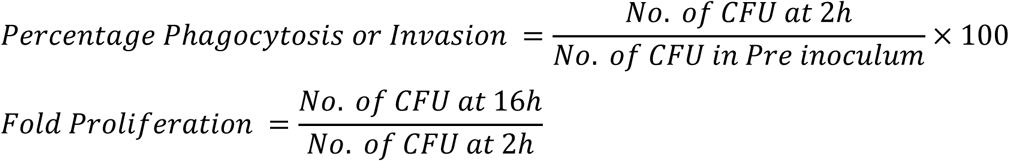

### RNA isolation and quantitative RT PCR

Bacterial strains were inoculated in LB broth, followed by subculture in LB broth at a ratio of 1:100. Bacterial cells were pelleted and lysed in TRIzol (RNAiso plus Takara). Total RNA was isolated from the TRIzol lysed sample as per the manufacturer’s protocol. Quantification of RNA was performed in NanoDrop (Thermo-Fischer Scientific). The quality of isolated RNA was detected by performing gel electrophoresis with 2% agarose gel. 1μg of RNA was subjected to Turbo DNase (Thermo-Fischer Scientific) treatment at 37°C for 30 minutes, followed by the addition of EDTA and heat inactivation at 65°C for 10 mins. The mRNA was reverse transcribed to cDNA using oligo dT primer, random hexamers, buffer, dNTPs, and reverse transcriptase (Takara) as per the manufacturer’s protocol. The expression profile of target genes was evaluated using specific primers by using TB green RT-qPCR master mix (Takara) in the Applied Biosystem QuantStudio-5 system. 16S rRNA and β-actin were used as internal controls for bacterial and host genes, respectively. All the reactions were set up in 384-well plates with three replicates for each sample. The primers used for RT-qPCR are listed in Table 2 [75], [76], [77].

### MTT Assay

Post 16h of infection, cells were treated with complete media containing 10μg/ml tetracycline for one hour to kill the intracellular bacteria, to negate the influence of MTT (3-(4,5-dimethylthiazol-2-yl)-2,5-diphenyltetrazolium bromide) metabolisation by the bacteria [78]. On completion of the tetracycline treatment, fresh incomplete media (i.e. DMEM only) containing 0.5mg/ml MTT (Sigma) was added to cells, followed by 3h of incubation at 37°C and 5% CO2 in the dark. After incubation, media is removed, and DMSO is added, followed by proper aspiration to dissolve the formazan dye. The optical density was measured at 590nm.

### Intracellular Reactive oxygen species (ROS) determination using H2DCFDA staining

Overnight cultures were inoculated into fresh LB media. Upon reaching an optical density of 0.1 (logarithmic phase), H2DCFDA was added at a final concentration of 10µM to 10^8^ CFU/ml of each strain in 1xPBS (pH 7.2) and incubated at 37°C for 30 minutes. Subsequently, 10^7^ H2DCFDA-stained bacterial cells were suspended in 1xPBS (pH 7.2) containing varying concentrations of hydrogen peroxide (1-5mM) and incubated at 37°C with orbital shaking for 2 hours. The samples were then transferred to a 96-well ELISA plate, and fluorescence was quantified using a Tecan-ELISA plate reader. (Ex-490nm/ Em-520nm).

### Bacterial nitric oxide (NO) determination using DAF-2DA staining

The production of nitric oxide (NO) indicates nitrosative stress. We evaluated the NO generation of the strains to *in vitro* nitrite stress by staining with DAF2-DA (4, 5- diamino fluorescein diacetate). DAF2-DA is a cell membrane-permeable dye that forms a membrane-impermeable adduct emitting fluorescence in NO presence. To mimic nitrite stress *in vitro*, 10^8^ CFU/ml of bacteria were treated with a range (500μM-5mM) of acidified nitrite (NaNO2 in PBS of pH 5.4) and incubated for 12 hours at 37°C. Post incubation, the cells are treated with 5μM DAF-DA and incubated for 30 minutes at 37°C. BD FACSverse flow cytometer was used to acquire the samples (Ex-491nm/ Em-513nm).

### Resazurin assay

Resazurin dye was used to evaluate the viability of bacteria when subjected to *in vitro* peroxide stress, acidified nitrite and peroxynitrite stress. 10^8^ CFU/ml of bacteria were treated with varying concentrations of H2O2 (1-5mM) and acidified nitrite (500μM-5mM), simulating oxidative and nitrosative stress, respectively. For peroxynitrite stress, a combination of H2O2 and acidified nitrite was used in a combined concentration range of 62.5μM -1mM. The treated bacteria were incubated for 12 hours at 37°C. On completion of treatment, resazurin was added to the bacteria at a final concentration of 1μg/ml, followed by an incubation at 37°C, for 2 hours. Post incubation, fluorescence intensity was measured in TECAN plate reader (Ex-540nm/ Em-590nm). Percentage viability was calculated by the formula:

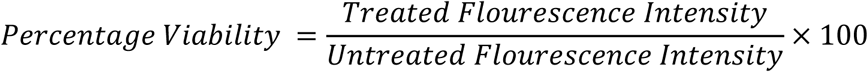

### Griess assay for estimating extracellular NO production by macrophages on infection

Griess assay detects the presence of nitrite, a stable product of NO breakdown. The protocol for the assay was adapted from [27], [79]. Briefly, 50μl of culture supernatants of infected macrophages or sodium nitrite standards (0.6, 1.2, 2.4, 4.8, 9.7, 19.5, 39, 78.1 and 156.2 μM) were treated with 50μl of 1% sulphanilamide (prepared in 5% phosphoric acid) and 50μl of 0.1% NED (N-1-naphthyl ethylene diamine dihydrochloride), followed by an incubation for 10 minutes at room temperature in the dark. On the appearance of a pink-purple colour product absorbance was measured at 545 nm.

### Measurement of NO production in infected macrophages

Intracellular NO production by RAW264.7 cells upon infection was measured by staining with DAF-2DA. 16h post-infection cells were treated with 5μM DAF-2DA and incubated for 30 minutes at 37°C. The cells were washed with 1xPBS and acquired in BD FACSverse flow cytometer (Ex-491nm/ Em-513nm).

### Immunofluorescence microscopy for nitrotyrosine residues, NOD1 and NOD2

RAW264.7 cells were seeded on coverslips and infected with eGFP-tagged strains [80]. The cells were fixed with 3.5% paraformaldehyde (PFA) 16h post-infection. The sample was stained with the rabbit-raised anti-nitrotyrosine or goat-raised anti-NOD1 or NOD2 antibody at a dilution of 1:100 (in 2.5% BSA, 0.01% saponin) overnight at 4°C, followed by staining with the corresponding secondary antibody tagged with a fluorophore at a dilution of 1:200 in 2.5% BSA, 0.01% saponin for one hour at room temperature. The antibodies used in this study are listed in Table 3. The coverslips were mounted on glass slides and sealed with transparent nail polish. The coverslips were imaged by using a Zeiss 880 microscope. Zeiss ZEN Black 2012 software was used to analyse the images. The formula used for calculating corrected total cell fluorescence (CTCF) is as follows:

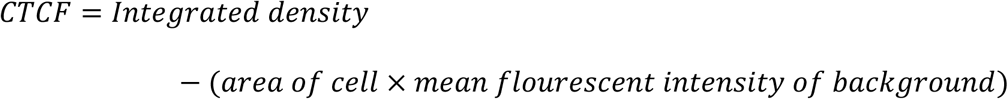

**Table 3.**
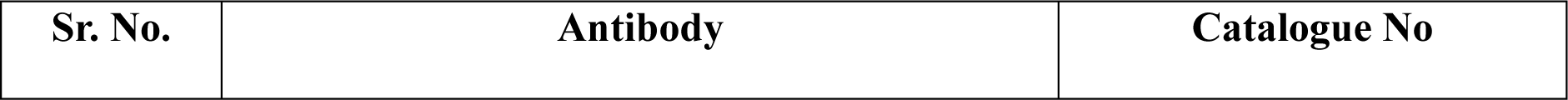
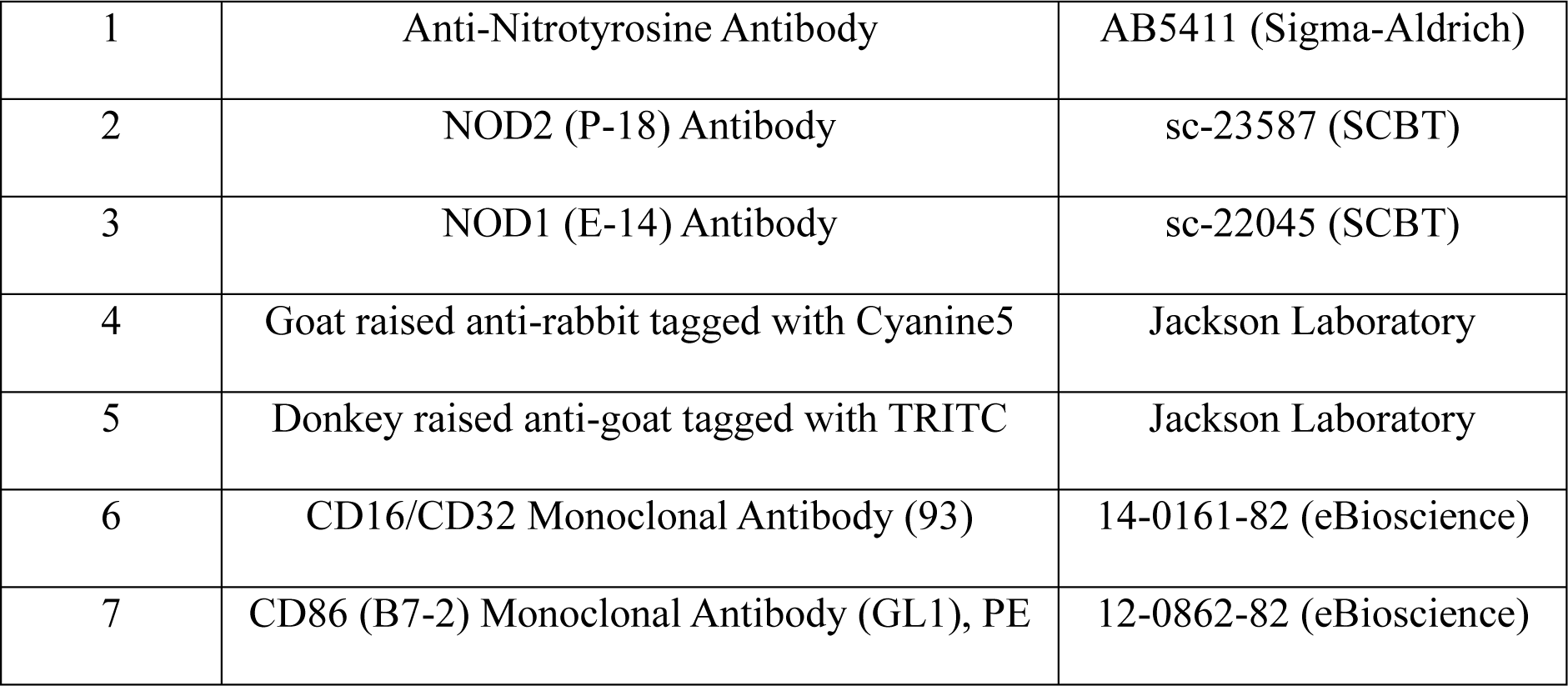
List of antibodies

### ELISA for estimating Nitrotyrosine and TNF-α

For the estimation of Nitrotyrosine, 16h post-infection, RAW264.7 cells were washed with 1x PBS and pelleted at 500g for 5 minutes. The cells were lysed with native lysis buffer (150 mM NaCl, 2 mM EDTA, 20 mM Tris, pH 8, 1% NP-40) at 4°C, with intermittent vortexing for 30 minutes, followed by centrifugation at 13,000 RPM for 15 minutes at 4°C. The protein concentration of the supernatant was estimated using the Bradford Reagent (Bio-Rad). Equal volumes of the supernatant, normalised to protein content, were applied to the pre-coated ELISA plates of the Nitrotyrosine ELISA kit (Abcam, ab210603), and the ELISA was performed according to the manufacturer’s protocol. To evaluate levels of TNF-α, the spent medium was collected from infected RAW264.7 cells 16 hours post-infection and subjected to centrifugation at 13,000 RPM, 4°C, for 10 minutes. The supernatant was added to the pre-coated TNF-α ELISA plates of the GENLISA Mouse TNF-α ELISA kit (Krishgen BioSystems, KB2145), and the ELISA was performed according to the manufacturer’s protocol. The final absorbance was measured at 450 nm wavelength using a Tecan Plate reader, and the concentrations of Nitrotyrosine and TNF-α were interpolated from a standard curve.

### Flow cytometry for assessing macrophage cell surface markers

The protocol was adapted from [81] and is as follows: Infected RAW 264.7 cells were washed and harvested in 1x PBS, 16 hours post-infection. Cells were centrifuged at 500g at 4°C, and the pellet was resuspended in Fc blocking solution, comprising anti-mouse CD16/CD32 antibody at a dilution of 1:200 in FACS buffer (1% BSA, 2 mM EDTA in 1x PBS), followed by a 30-minute incubation on ice. Post-incubation, the cells were washed and stained with a PE-conjugated CD86 antibody at a dilution of 1:200 for 1 hour on ice. Subsequently, the cells were washed and fixed with 2% PFA in PBS for 10 minutes. The cells were washed with 1x PBS, resuspended in FACS buffer, and acquired using CytoFLEX flow cytometer by Beckman Coulter Life Sciences. The data were then analysed in CytExpert software.

### Bacterial burden in murine model

Specific pathogen-free (SPF) 6-8 weeks old C57BL/6 (*iNOS^+/+^*) and *iNos^-/-^* (*iNOS* knockout in C57BL/6 background) mice were segregated in cohorts of n=5 or 6. For oral infection, C57BL/6 mice were orally gavaged with 10^7^ CFU of STM WT or STM*ΔuppP*. For *iNOS^-/-^*mice, a dose of 10^5^ CFU was administered orally. The infected mice were ethically euthanised 5 days post-infection. For intraperitoneal injection, C57BL/6 (*iNOS^+/+^*) and *iNOS^-/-^* mice were injected with a dose of 10^4^ CFU, followed by ethical euthanasia 3 days post-infection. Bacterial burden was enumerated in the homogenised lysate of organs by serial dilution plating on *Salmonella*-*Shigella* agar.

### Mouse survival assay

Specific pathogen-free (SPF) 6-8 weeks C57BL/6 mice were orally gavaged with 10^7^ CFU of STM WT or STM*ΔuppP* (n=6) and monitored daily for mortality. The data is represented as survival percentage.

### Statistical analysis

Data was analysed and graphed using the GraphPad Prism 8 software (San Diego, CA). Statistical significance was determined by Student’s t-test, One-way ANOVA or Two-way ANOVA to obtain p values. Adjusted p-values below 0.05 are considered statistically significant. The results are expressed as mean ± SD unless mentioned otherwise.

## Ethics statement

All the animal experiments have been approved by the Institutional Animal Ethics Clearance Committee (IAEC), Indian Institute of Science, Bangalore. The registration number is 48/1999/CPCSEA. The guidelines formulated by the Committee for the Purpose of Control and Supervision of Experiments on Animals (CPCSEA) were strictly followed. The ethical clearance number for this study is CAF/Ethics/853/2021.

## Acknowledgements

We acknowledge the Real-Time Facility, Departmental Confocal Microscopy Facility and Central Instrumentation Facility, at the Department of Microbiology and Cell Biology (MCB), the Division of Biological Sciences Flow Cytometry Facility, the Atomic Force Microscopy (AFM) Facility at the Department of Bioengineering (BE), Advanced Facility for Microscopy and Microanalysis (AFMM) and Central Animal Facility at the Indian Institute of Science (IISc). Mrs. Deepika and Mrs. Monisha are acknowledged for their help in acquiring Scanning Electron Microscopy (SEM) images and AFM images, respectively. We are grateful to Mr. Munish and Mrs. Navya for helping with the data acquisition for flow cytometry. We thank Mr. Arjun Suresh for his assistance with confocal microscopy image acquisition. Dr Abhilash Vijay Nair is acknowledged for providing us with the gene of interest for our study.

## Author Contributions

Conceptualisation: RV, DM, KP, and DC; Methodology: RV, DM, KP, KCN (PG isolation and analysis); Investigation: RV, DM, KP, KCN (PG isolation and analysis); Data curation: RV, Formal analysis: RV, DM, KP, Fund acquisition: DC, MR (PG isolation and analysis) Project administration: RV and DC, Writing-original draft: RV, Editing and proofreading: RV, DM, KP, KCN, MR and DC, Supervision: DC.

## Supplementary Figures

**Figure S1.**
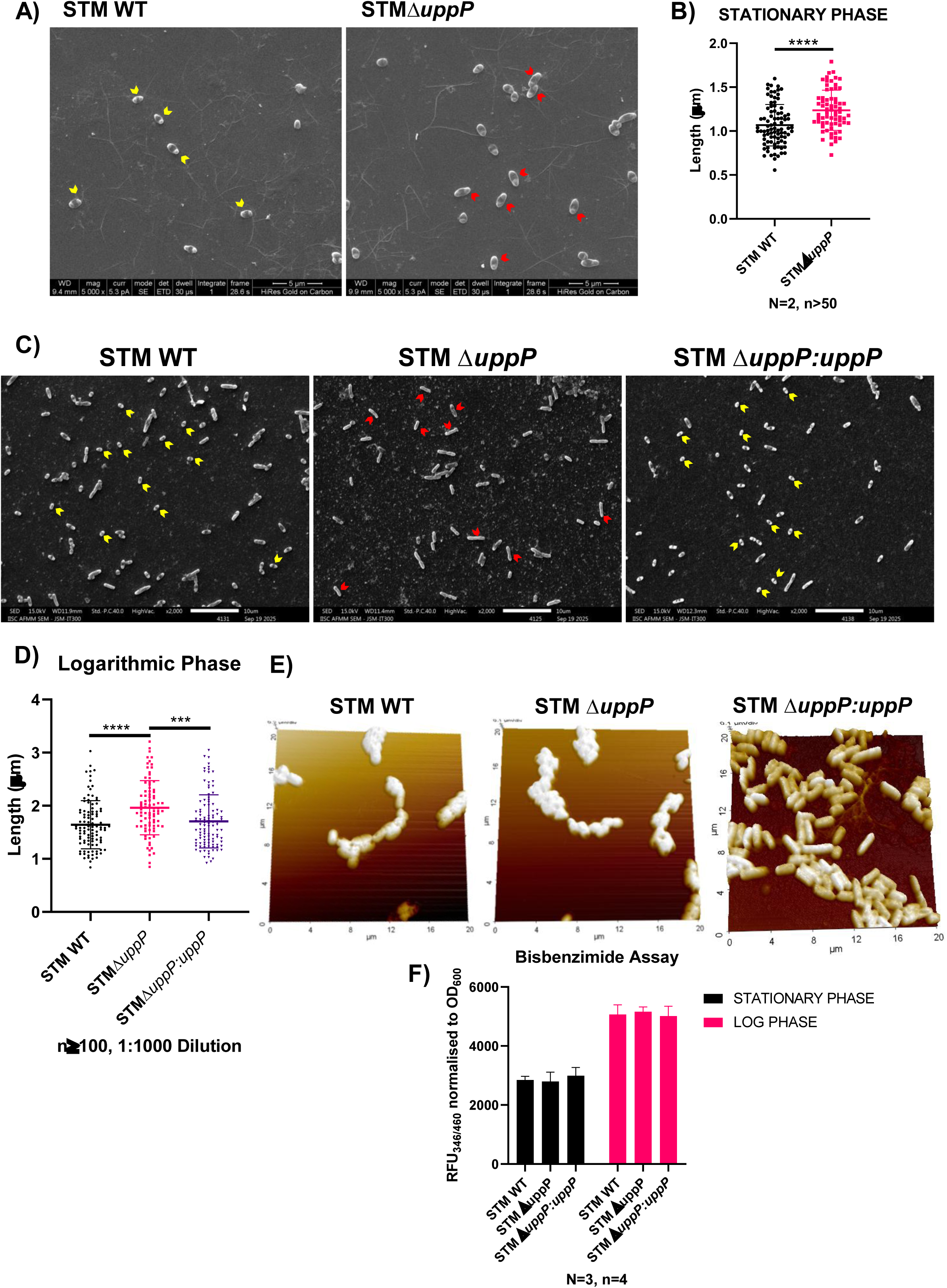
Loss of *uppP* affects the bacterial cell dimensions A) Representative scanning electron micrographs of STM WT and STM *ΔuppP* stationary phase cells. Red and yellow arrows represent cells with increased and decreased length, respectively. Scale bar = 5μm B) Quantification of the length of the stationary phase bacterial cells captured in the SEM images (A). Data are representative of N=2, n≥50 ± SD (standard deviation). An unpaired two-tailed Student’s t-test was performed to obtain the p-values. (****P ≤ 0.0001, ***P ≤ 0.001, **P ≤ 0.01, *P ≤ 0.05 ns- non-significant) C) Representative scanning electron micrographs of STM WT, STM *ΔuppP* and STM *ΔuppP*:*uppP* logarithmic phase cells (1:1000 dilution). Red and yellow arrows represent cells with increased and decreased length, respectively. Scale bar = 5μm D) Quantification of the length of the logarithmic phase (1:1000 dilution) bacterial cells captured in the SEM images (C). Data are representative of N=2, n≥50 ± SD (standard deviation). One-way ANOVA was performed to obtain the p-values. (****P ≤ 0.0001, ***P ≤ 0.001, **P ≤ 0.01, *P ≤ 0.05 ns- non-significant) E) Representative atomic force micrographs of STM WT, STM *ΔuppP* and STM *ΔuppP*:*uppP*. A 20 x 20μm^2^ area from each coverslip was selected for imaging. F) Measurement of outer membrane porosity of STM WT, STM *ΔuppP* and STM *ΔuppP*:*uppP*, in the stationary and logarithmic phase by bisbenzimide uptake assay. Data are representative of N=3, n=4 ± SD (standard deviation). Two-way ANOVA was performed to obtain the p-values. (****P ≤ 0.0001, ***P ≤ 0.001, **P ≤ 0.01, *P ≤ 0.05 ns- non-significant)

**Figure S2.**
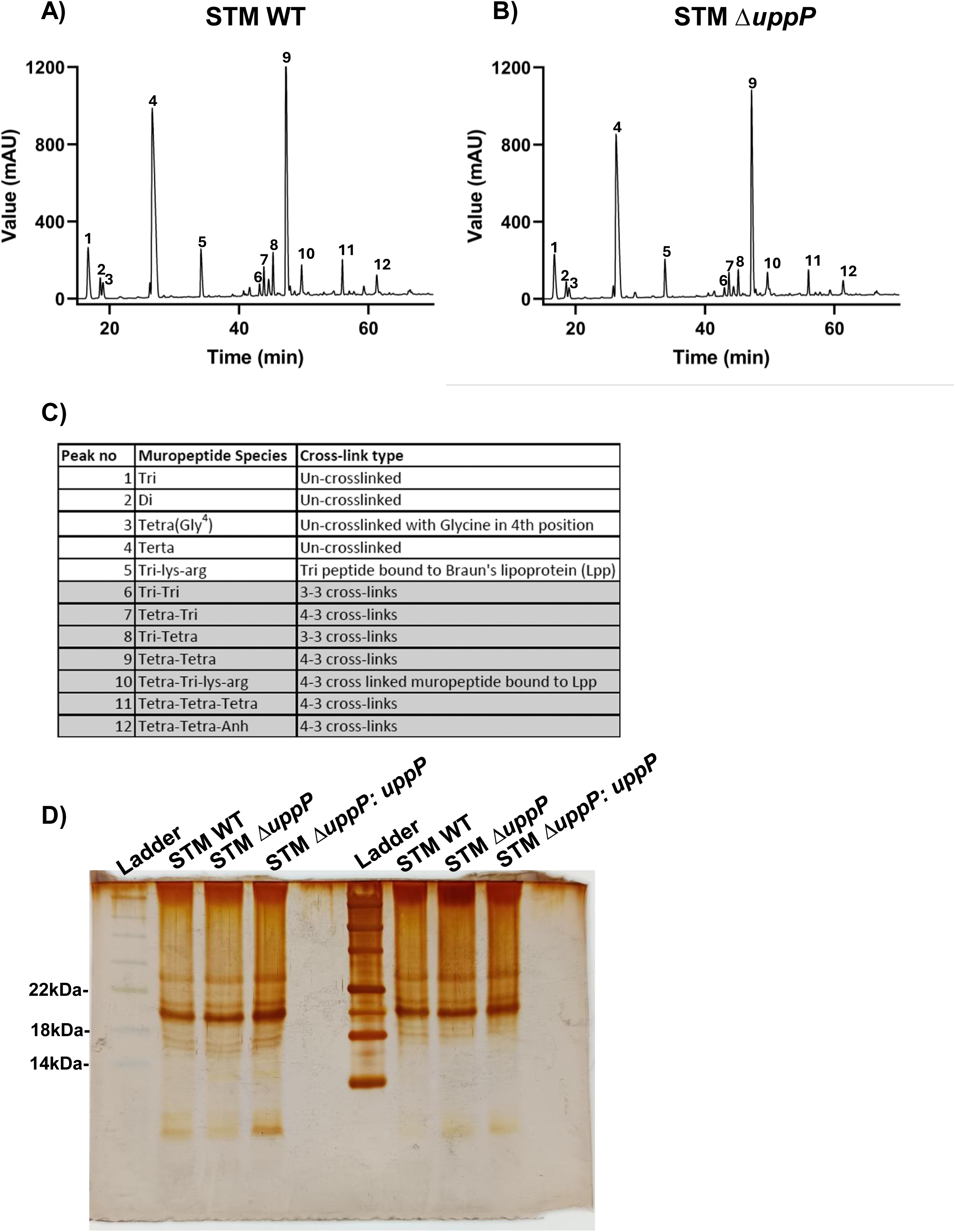
Peptidoglycan and LPS composition of STM Δ*uppP* A-B) HPLC chromatograms showing petidoglycan composition of STM WT and STM *ΔuppP* strains grown in Luria Bertani broth. Strains were grown till they reached the early logarithmic phase (OD600nm ∼1). Data are representative of N=2. The identities of the peaks, according to their mass, are represented in the table (C). C) Table summarising the identity of the muropeptide fragment peaks observed in HPLC chromatograms (A) and (B). D) Silver staining gel of LPS isolated from STM WT, STM *ΔuppP* and STM *ΔuppP*:*uppP*. Data are representative of N=2.

**Figure S3.**
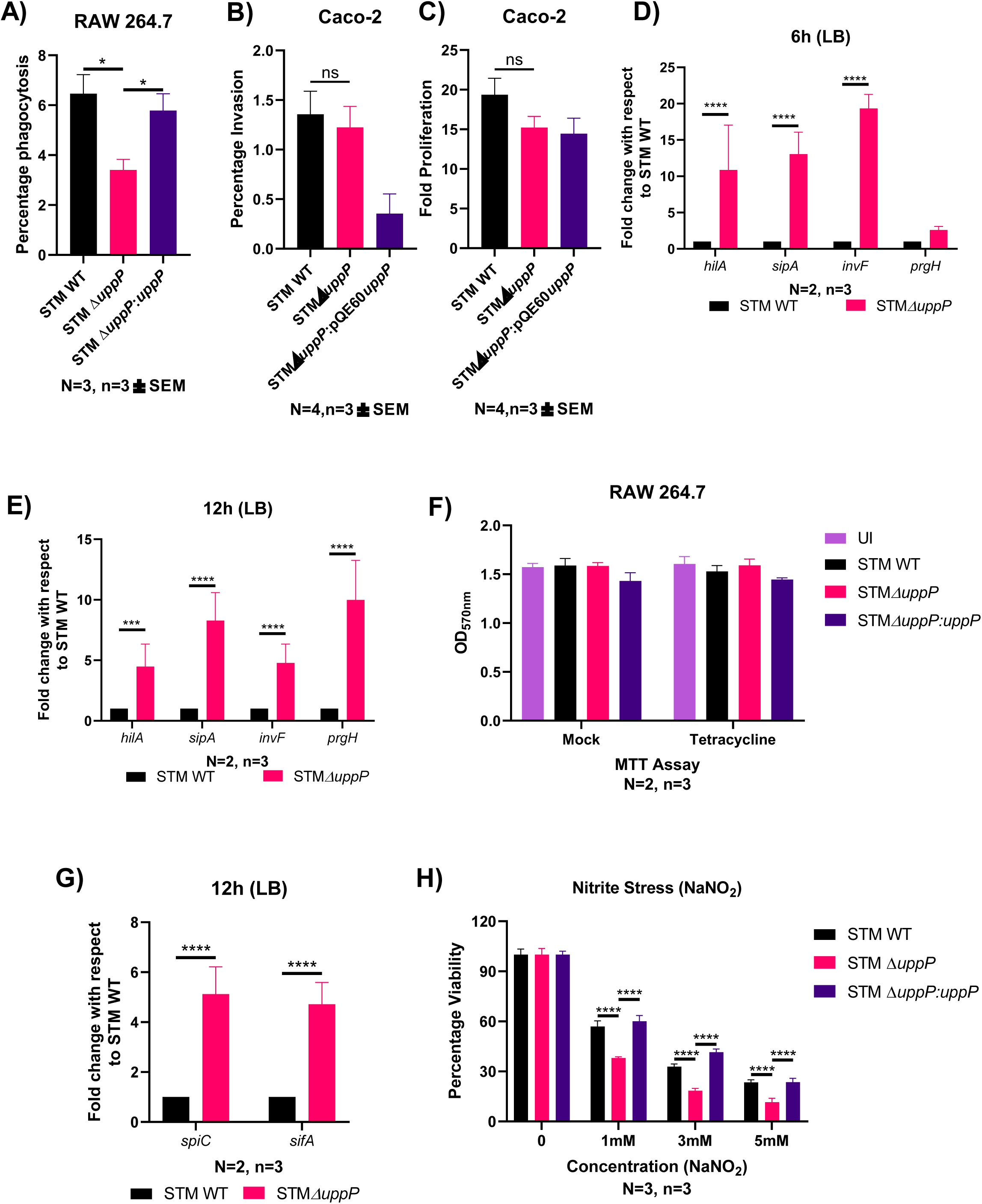
*uppP* facilitates the intracellular survival of *Salmonella* in macrophages A) Percentage phagocytosis of STM WT, STM *ΔuppP*, and STM *ΔuppP:uppP*, within RAW 264.7 macrophages (MOI=10). Results are mean of N=3, n=3 ± SEM (standard error of mean). One-way ANOVA was performed to obtain the p-values. (****P ≤ 0.0001, ***P ≤ 0.001, **P ≤ 0.01, *P ≤ 0.05 ns- non-significant) B-C) Percentage invasion (B) and fold proliferation (C) of STM WT, STM *ΔuppP* and STM *ΔuppP:uppP* within colorectal adenocarcinoma cell line Caco-2 (MOI=10). Results are mean of N=3, n=3 ± SEM (standard error of mean). An unpaired two-tailed student’s t-test was performed to obtain the p-values. (****P ≤ 0.0001, ***P ≤ 0.001, **P ≤ 0.01, *P ≤ 0.05 ns-non-significant) D-E) Quantification of the expression of *hilA*, *prgH*, *invF* and *sipA* during logarithmic (E) and stationary (F) phase in STM WT and STM Δ*uppP* cultured in LB broth. Results are representative of N=2, n=3 ± SD. Two-way ANOVA was performed to obtain the p-values. (****P ≤ 0.0001, ***P ≤ 0.001, **P ≤ 0.01, *P ≤ 0.05 ns- non-significant) F) Cell viability assay of RAW 264.7 cells, 16h post-infection with STM WT, STM *ΔuppP*, and STM *ΔuppP:uppP*. Results are representative of N=2, n=3 ± SD. Two-way ANOVA was performed to obtain the p-values. (****P ≤ 0.0001, ***P ≤ 0.001, **P ≤ 0.01, *P ≤ 0.05 ns-non-significant) G) Quantification of the expression of *spiC* and *sifA* during the stationary phase in STM WT and STM Δ*uppP* cultured in LB broth. Results are representative of N=2, n=3 ± SD. Two-way ANOVA was performed to obtain the p-values. (****P ≤ 0.0001, ***P ≤ 0.001, **P ≤ 0.01, *P ≤ 0.05 ns- non-significant) H) Percentage viability of resazurin assay of STM WT, STM Δ*uppP* and STM *ΔuppP:uppP* in the presence of varying concentrations of NaNO2 (*in vitro* nitrite stress). Results are representative of N=3, n=3 ± SD. Two-way ANOVA was performed to obtain the p-values. (****P ≤ 0.0001, ***P ≤ 0.001, **P ≤ 0.01, *P ≤ 0.05 ns- non-significant)

**Figure S4.**
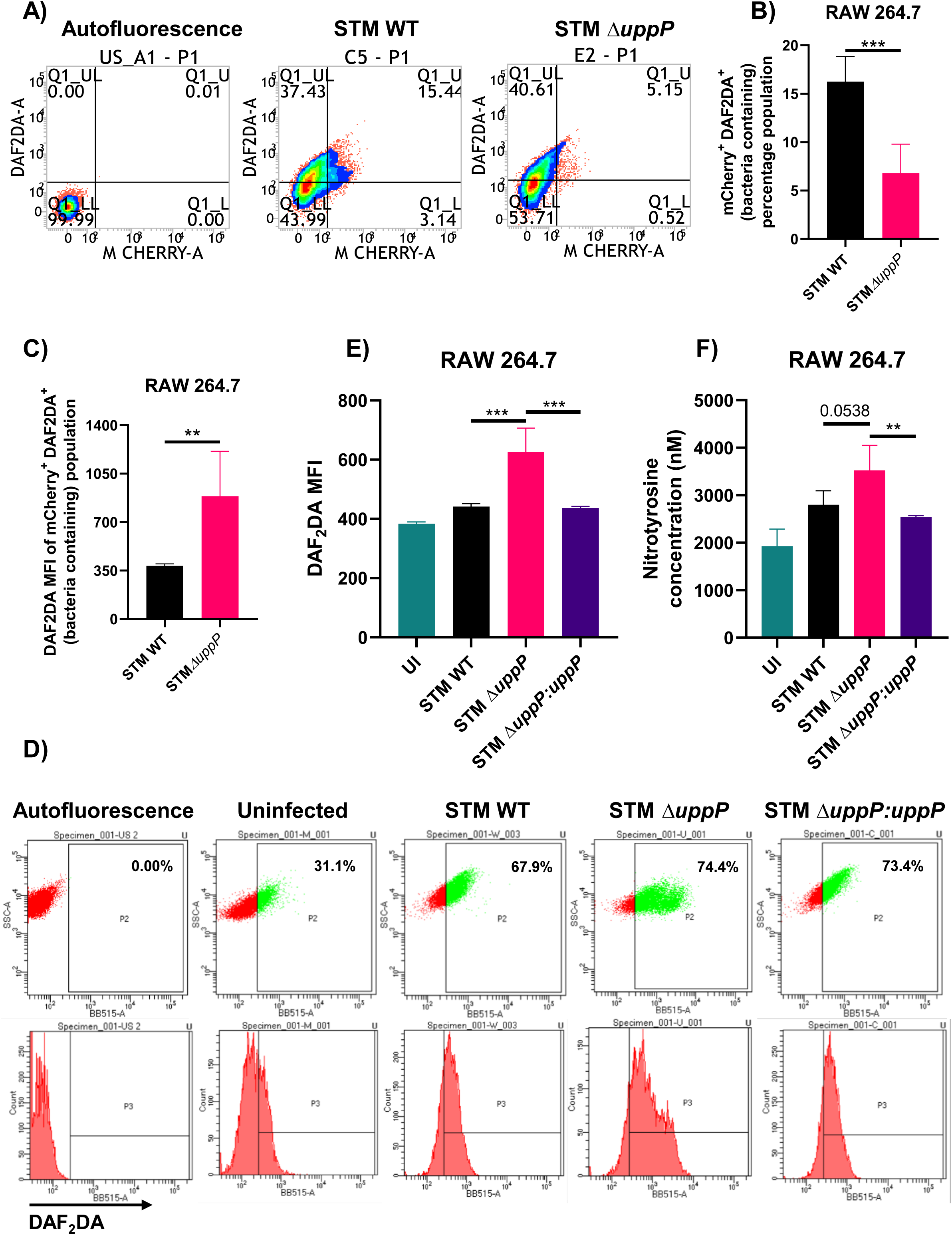
STM Δ*uppP* upregulates nitric oxide (NO) levels in RAW 264.7 cells A-C) Alternative gating strategy for Fig. 3A-B. Estimation of NO in macrophages infected with mCherry-tagged STM WT and STM Δ*uppP* by DAF2DA staining followed by flow cytometry. Representative DAF2DA vs mCherry scatter plots of STM WT and STM Δ*uppP* (A). Quantification of mCherry- DAF2DA double-positive percentage population (B) and DAF2DA median fluorescence intensity (MFI) (C). Results are representative of N=3, n=4 ± SD. Unpaired two-tailed Student’s t-test was performed to obtain the p-values. (****P ≤ 0.0001, ***P ≤ 0.001, **P ≤ 0.01, *P ≤ 0.05 ns- non-significant) D-E) Estimation of NO, 16h post-infection with STM WT, STM Δ*uppP* and STM Δ*uppP:uppP* in RAW264.7 cells by DAF2DA staining followed by flow cytometry. Representative SSC-A vs DAF2DA dot plots and Count vs DAF2DA histogram plots of STM WT, STM Δ*uppP*, and STM Δ*uppP:uppP* infected RAW 264.7 cells (D). Quantification of median fluorescence intensity (MFI) of DAF2DA-positive population (E). Results are representative of N=2, n=4 ± SD. One-way ANOVA was performed to obtain the p-values. (****P ≤ 0.0001, ***P ≤ 0.001, **P ≤ 0.01, *P ≤ 0.05 ns- non-significant) F) Estimation of nitrotyrosine through ELISA, in cell lysates of RAW 264.7 cells infected with STM WT, STM *ΔuppP*, and STM Δ*uppP:uppP*. Results are representative of N=2, n=2 ± SD. One-way ANOVA was performed to obtain the p-values. (****P ≤ 0.0001, ***P ≤ 0.001, **P ≤ 0.01, *P ≤ 0.05 ns- non-significant)

**Figure S5.**
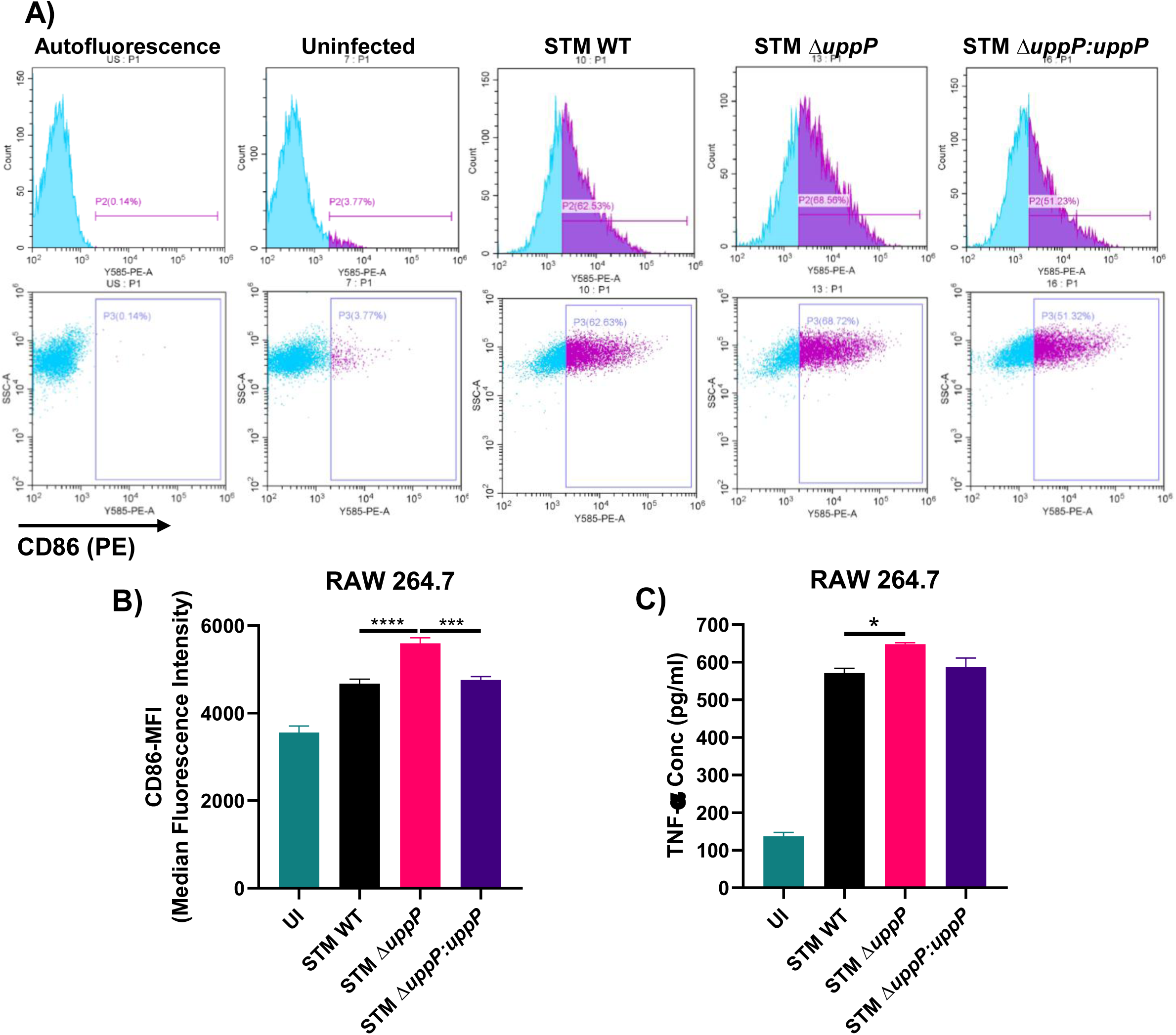
STM Δ*uppP* triggers an enhanced immune response in RAW 264.7 cells A-B) Flow cytometric analysis of cell surface levels of CD86 in macrophages, 16h post-infection with STM WT, STM Δ*uppP* and STM Δ*uppP:uppP*. Representative Count vs CD86 (PE) histogram plots and SSC-A vs CD86 (PE) dot plots of STM WT, STM Δ*uppP* and STM Δ*uppP:uppP* infected RAW 264.7 cells (A). Quantification of median fluorescence intensity (MFI) of CD86 (PE) positive population (B). Results are representative of N=2, n=4 ± SD. One-way ANOVA was performed to obtain the p-values. (****P ≤ 0.0001, ***P ≤ 0.001, **P ≤ 0.01, *P ≤ 0.05 ns- non-significant) C) Estimation of pro-inflammatory cytokine TNF-α production through ELISA in cell supernatants of RAW 264.7 macrophages infected with STM WT, STM Δ*uppP*, and STM Δ*uppP:uppP* for 16 hours. Results are mean of N=2, n=2 ± SEM (standard error of mean). One-way ANOVA was performed to obtain the p-values. (****P ≤ 0.0001, ***P ≤ 0.001, **P ≤ 0.01, *P ≤ 0.05 ns- non-significant)

**Figure S6.**
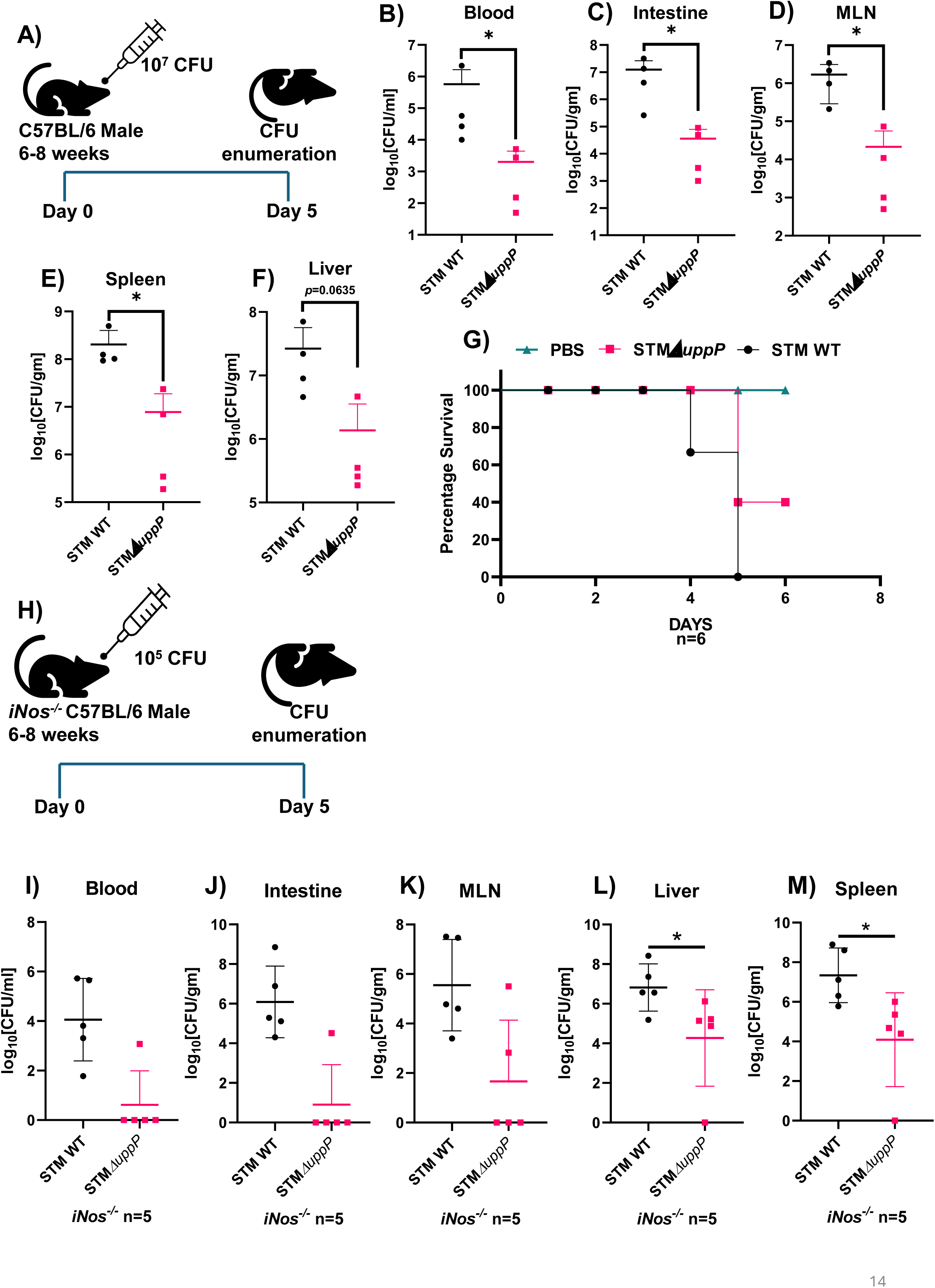
UppP facilitates *Salmonella* survival in the *in vivo* murine model A) Schematic representation of the experimental protocol followed for enumerating bacterial burden in tissues upon oral gavage in C57BL/6 (*iNOS^+/+^*) mice. B-F) Enumeration of bacterial burden in the blood (B), intestine (C), MLN (D), spleen (E) and liver (F) on infection via oral route in C57BL/6 (*iNOS^+/+^*) mice. Results are representative of N=2, n=6 ± SD (standard deviation). Mann-Whitney test was performed to obtain the p-values. (****P ≤ 0.0001, ***P ≤ 0.001, **P ≤ 0.01, *P ≤ 0.05 ns- non-significant) G) Percentage survival of C57BL/6 (*iNOS^+/+^*) mice infected with STM WT and STM*ΔuppP* via the oral route. H) Schematic representation of the experimental protocol followed for enumerating bacterial burden in tissues upon oral gavage in *iNOS^-/-^* mice. I-M) Enumeration of bacterial burden in the blood (I), intestine (J), MLN (K), liver (L) and spleen (M) on infection via oral route in *iNOS^-/-^* mice. Results are representative of n=5 ± SD (standard deviation). Mann-Whitney test was performed to obtain the p-values. (****P ≤ 0.0001, ***P ≤ 0.001, **P ≤ 0.01, *P ≤ 0.05 ns- non-significant)

**Figure S7.**
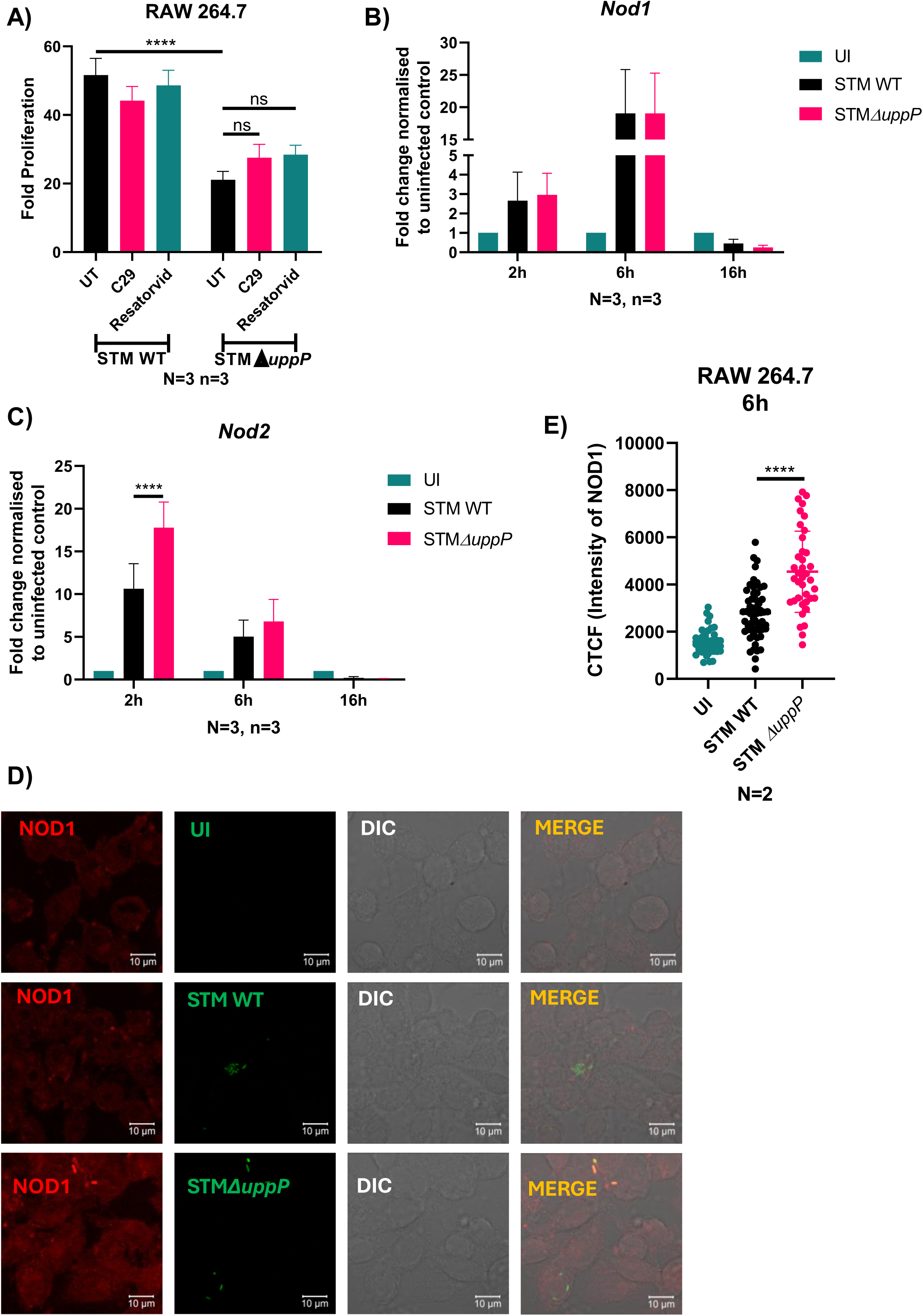
NOD receptors contribute to the attenuated survival of STM Δ*uppP* in RAW 264.7 cells A) Fold proliferation of STM WT and STM *ΔuppP* within RAW 264.7 macrophages on treatment with C29 and Resatorvid, MOI=10. Results are mean of N=3, n=3 ± SEM (standard error of mean). One-way ANOVA was performed to obtain the p-values. (****P ≤ 0.0001, ***P ≤ 0.001, **P ≤ 0.01, *P ≤ 0.05 ns- non-significant) B-C) Quantification of mRNA expression of *Nod1* (B) and *Nod2* (C) in RAW264.7 cells infected with STM WT and STM Δ*uppP* for 2h, 6h and 16h. Results are representative of N=3, n=3 ± SD. Two-way ANOVA was performed to obtain the p-values. (****P ≤ 0.0001, ***P ≤ 0.001, **P ≤ 0.01, *P ≤ 0.05 ns- non-significant) D-E) Representative immunofluorescence images of RAW 264.7 cells infected with STM WT and STM *ΔuppP* for 6h, MOI=10, immunostained for NOD1 (D). Quantification of corrected total cell fluorescence (CTCF) of NOD1 in infected cells (E). Results are representative of N=2, n≥50 ± SD. One-way ANOVA was performed to obtain the p-values. (****P ≤ 0.0001, ***P ≤ 0.001, **P ≤ 0.01, *P ≤ 0.05 ns- non-significant)

